# A binding site for phosphoinositides described by multiscale simulations explains their modulation of voltage gated sodium channels

**DOI:** 10.1101/2023.07.16.547149

**Authors:** Yiechang Lin, Elaine Tao, James P Champion, Ben Corry

## Abstract

Voltage gated sodium channels (Na_v_) are membrane proteins which open to facilitate the inward flux of sodium ions into excitable cells. In response to stimuli, Na_v_ channels transition from the resting, closed state to an open, conductive state, before rapidly inactivating. Dysregulation of this functional cycle due to mutations causes diseases including epilepsy, pain conditions and cardiac disorders, making Na_v_ channels a significant pharmacological target. Phosphoinositides are important lipid cofactors for ion channel function. The phosphoinositide PI(4,5)P_2_ decreases Na_v_1.4 activity by increasing the difficulty of channel opening, accelerating fast inactivation and slowing recovery from fast inactivation. Using multiscale molecular dynamics simulations, we show that PI(4,5)P_2_ binds stably to inactivated Na_v_ at a conserved site within the DIV S4-S5 linker, which couples the voltage sensing domain (VSD) to the pore. As the Nav C-terminal domain is proposed to also bind here during recovery from inactivation, we hypothesise that PI(4,5)P_2_ prolongs inactivation by competitively binding to this site. In atomistic simulations, PI(4,5)P_2_ reduces the mobility of both the DIV S4-S5 linker and the DIII-IV linker, responsible for fast inactivation, slowing the conformational changes required for the channel to recover to the resting state. We further show that in a resting state Na_v_ model, phosphoinositides bind to VSD gating charges, which may anchor them and impede VSD activation. Our results provide a mechanism by which phosphoinositides alter the voltage dependence of activation and the rate of recovery from inactivation, an important step for the development of novel therapies to treat Na_v_-related diseases.

**Significance:** Voltage-gated sodium channels form pores in the membrane to mediate electrical activity in nerve and muscle cells. They play critical roles throughout the human body and their dysfunction leads to diseases including epilepsy, cardiac arrhythmias and pain disorders. Membrane lipids called phosphoinositides have recently been shown to reduce the activity of a voltage-gated sodium channel, but the molecular basis of this mechanism is not known. Here we use simulations to reveal where these lipids bind to the channels and how they reduce channel activity by making it harder for the pores to open and slower to subsequently recover to the closed resting state.

## Introduction

Voltage-gated sodium (Na_v_) channels are critical to the regulation of brain activity, cardiac rhythm, and muscle contraction. Expressed in the membranes of excitable cells, Na_v_ channels respond to membrane depolarization to open a pore that facilitates the selective flow of sodium current into the cell, initiating the action potential. In mammals, the Na_v_ channel family consists of nine subtypes (Na_v_ 1.1-1.9), distributed throughout the central and peripheral nervous system, as well as in cardiac and skeletal muscle (1). The Na_v_1.4 subtype is predominantly expressed in skeletal myofibers, where it initiates muscle contraction. Genetic mutations in this subtype are associated with various motor dysfunctions, such as both hyperkalaemic and hypokalaemic periodic paralyses (2-4). Na_v_1.7 is found in peripheral sensory neurons and is responsible for nociception. Several pain disorders, such as inherited erythromelalgia and small fibre neuropathy arise from gain-of-function Na_v_1.7 mutations (5). Both Na_v_ subtypes have been investigated as promising pharmacological targets for the treatment of myopathy and pain conditions (5, 6).

Structurally, Na_v_ channels consist of four homologous domains (DI-DIV) arranged in a domain-swapped configuration (Fig. 1A-B). Each domain comprises of six transmembrane helices (S1-S6). The central pore domain is formed by S5, S6 helices and selectivity filter (SF), while the four peripheral voltage-sensing domains (VSDs) are formed by S1-S4 (7, 8). The pore domain also features lateral fenestrations that provide a pathway for the access of small molecules to the pore via the membrane (9) and have been shown in computational studies to be accessible to lipid tails (10-13). Additionally, the C-terminal domain (CTD) extends from the DIV S6 helix into the cytoplasm, where it is thought to associate with the DIII-IV and DIV S4-S5 linkers in the resting state (14).

**Fig. 1.**
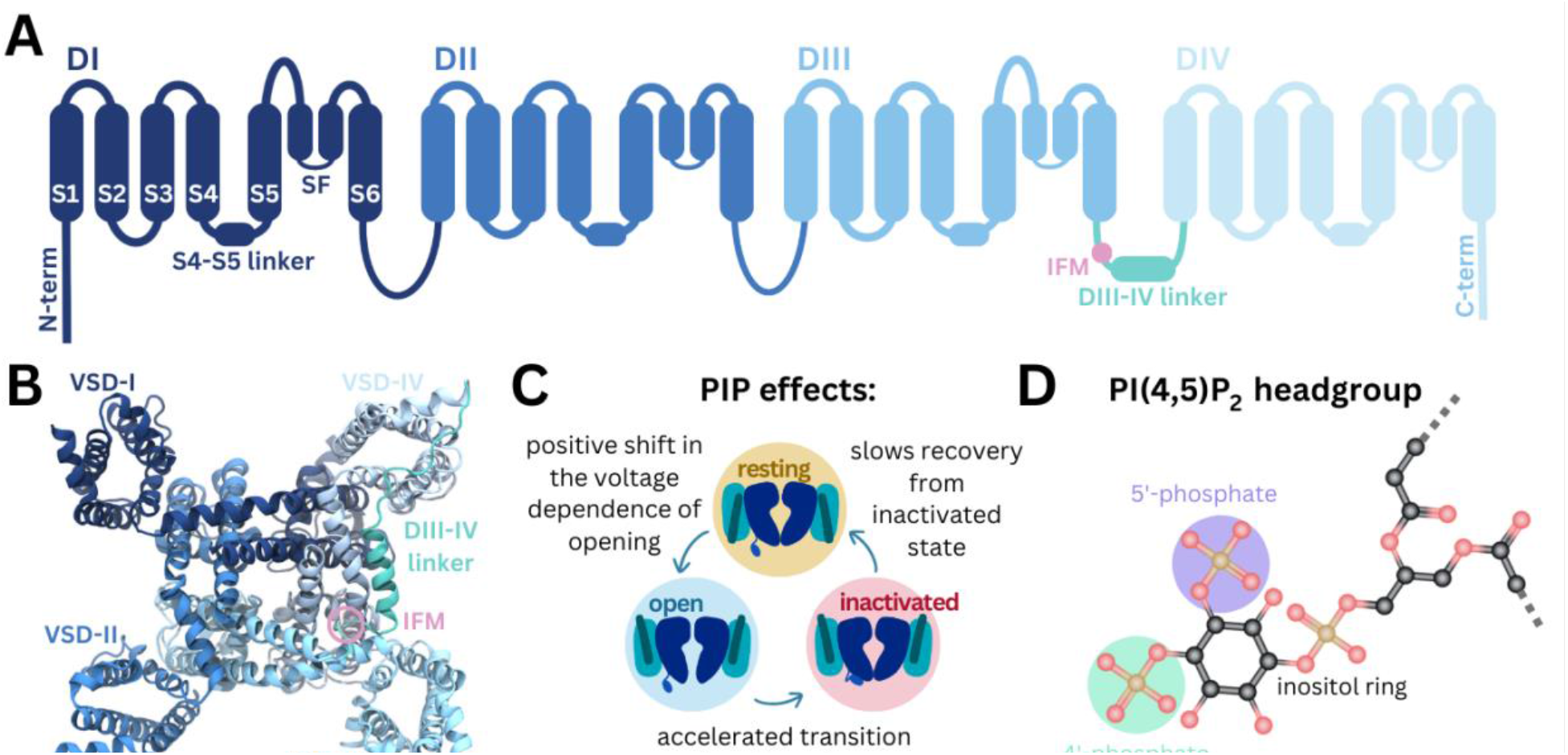
Structure of Na_v_ and modulation by phosphoinositides. **(A)** Na_v_ channel topology featuring transmembrane helices (S1-S6), the selectivity filter (SF) and the DIII-IV linker (containing the IFM motif) located between DIII and DIV. **(B)** Na_v_1.4 structure (6agf) showing the four domain-swapped voltage sensor domains (VSDs I-IV), pore and DIII-IV linker on the intracellular side. **(C)** Summary of PI(4,5)P_2_ effects on transitions between Na_v_ channel functional states (25) **(D)** Structure of the PI(4,5)P_2_ headgroup with the 4’- and 5’-phosphates indicated

Na_v_ channels adopt distinct functional states during the cycle of membrane depolarization and repolarization (Fig. 1C). In the resting state, the pore is closed and VSD S4 helices are in the down, deactivated state. Membrane depolarization triggers the asynchronous transition of VSD I-III S4 helices to the up, activated conformation, causing Na_v_ channels to open (15-17). During prolonged depolarization, VSD-IV moves upwards, causing the channels to adopt a fast inactivated state in which a three-residue Ile-Phe-Met (IFM) hydrophobic motif located on the intracellular linker between DIII and DIV (DIII-IV linker) allosterically closes the pore (18). The CTD is proposed to bind at the S4-S5 linker of VSD-IV and sequester the DIII-IV linker during the resting state. During fast inactivation, the CTD dissociates from VSD-IV and releases the DIII-IV linker to allow IFM binding (14). Upon repolarization, the VSDs deactivate, the IFM motif disassociates, the CTD rebinds to VSD-IV and the pore returns to the resting state.

Phosphoinositides (PIPs) are important cellular signaling molecules found on the cytoplasmic leaflet of the mammalian cell membrane. They can exist in seven forms, with phosphorylation possible at one (PIP1), two (PIP2) or all three (PIP3) positions on the inositol ring, at the 3’, 4’ and/or 5’ carbons. PIPs, particularly PI(4,5)P_2_, featuring phosphates at the 4’ and 5’ carbon positions (Fig. 1D), are known to bind and modulate the activity of numerous ion channels families (19). These include voltage-gated ion channels, some of which have been resolved with PI(4,5)P_2_ bound (20, 21). PIP is known to interact with the VSDs of different potassium channels, to stabilize the positive gating charges and support the voltage-sensing mechanism (22). By also binding at the VSD-pore interface in channels such as K_v_7.1, PIP is proposed to facilitate coupling of VSD movement to pore opening (20, 23, 24). PI(4,5)P_2_ also forms specific interactions with VSD-II of Ca_v_2.2 in the down-state, making channel activation more difficult (21). Although PI(4,5)P_2_ is known to bind to numerous voltage gated ion channels, its effects on channel gating and function are complex and yet to be fully elucidated.

Recent experiments show that Na_v_1.4 channel kinetics and voltage dependence are modulated by PI(4,5)P_2_ (25). PI(4,5)P_2_ inhibits Na_v_1.4 by causing a depolarizing shift in voltage dependence of activation, accelerating transition to the inactivated state and slowing recovery from inactivation, resulting in reduced peak current and suppression of late current (summarized in Fig. 1C). While this is likely to occur via a direct interaction with Na_v_1.4, the structural basis of PI(4,5)P_2_ modulation remains to be understood.

Here, we used a combination of coarse-grained and atomistic MD simulations to identify a putative PIP binding site to inactivated Na_v_1.4 in VSD-IV and the DIII-IV linker. We analyze the atomistic level interactions between the positively charged residues at this site with PI(4,5)P_2_ and PI(4)P, comparing this with structurally resolved PIP binding sites in the related ion channels Ca_v_2.2 and K_v_7.1. Consistent with the sequence conservation at the identified site, we find that PIPs also bind to Na_v_1.7 in coarse-grained simulations, with notable differences dependent on VSD conformation states. This work provides insight into how PIPs can negatively regulate Na_v_ channels, a first step for the potential development of PIP-analogue sodium channel inhibitors.

## Results

To investigate how diverse lipid species interact with Na_v_1.4, we carried out coarse-grained simulations of Na_v_1.4 (PDB ID: 6agf, inactivated state) embedded in a complex mammalian membrane for 16 μs in triplicate (Fig. 2A). Glycosphingolipid (GM), phosphoinositide (PIP) and diacylglycerol (DAG) were highly enriched around Na_v_1.4 (Fig. 2B). Additionally, we observed modest enrichment of lysophosphatidylcholine (LPC), phosphatidylinositol (PI), phosphatidylserine (PS) and phosphatidylethanolamine (PE) and slight depletion of ceramide (CER), sphingomyelin (SM), phosphatidylcholine (PC) and cholesterol (CL). To investigate specific interactions, we generated z-density maps and calculated the per-residue occupancy of the 12 different lipid types (Fig. 2C, Fig. S1-2). Binding residues of interest were identified by constructing occupancy distributions by residue for each lipid type and identifying outlying values with high occupancies (Fig. S3). DG lipids form significant interactions within the lateral fenestrations of Na_v_1.4 (Fig. S2). LPC and PI also frequently interact with different VSD residues (Fig. S1).

**Fig. 2.**
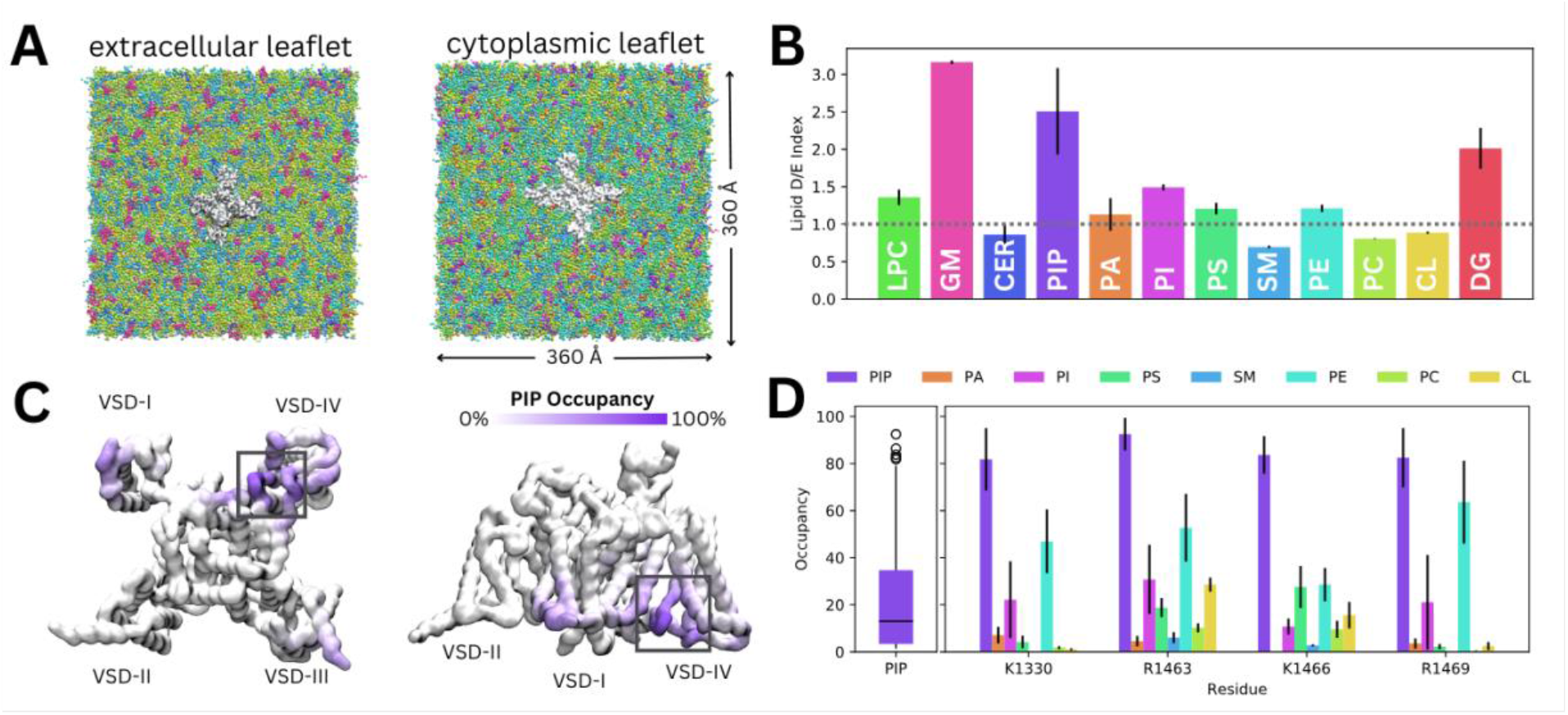
Lipid fingerprint and binding of all PIP types to Na_v_1.4. **(A)** Na_v_1.4 embedded in a 360 Å x 360 Å model mammalian membrane containing 63 lipid species. **(B)** Lipid depletion enrichment index of lipids around Na_v_1.4 grouped into 12 headgroup classes **(C)** Na_v_1.4, shown from the intracellular (left) and membrane (right) sides, colored by PIP occupancy (darker purple = greater PIP occupancy) **(D)** Distribution of PIP binding occupancies (left) and occupancy of lipid species at 4 residues with the highest PIP occupancy.

Given our interest in understanding the modulation of Na_v_1.4 by PIPs, we focus on their interactions for the remainder of this manuscript. Despite the very low concentration of PIPs (0.5% each of PIP1, PIP2 and PIP3) in the mammalian membrane, they are highly enriched around Na_v_1.4, particularly near the DIII-IV linker and the VSD-IV (Fig. 2C). Contact analysis revealed a putative PIP binding site involving the K1330 residue of the DIII-IV linker and residues R1463, K1466 and R1469 in the DIV S4-S5 linker, which connects the pore and VSD-IV (Fig. 2D). Across all three replicates, only residues within this site were occupied by PIPs for more than 80% (Quartile 3 + 1.5 x Interquartile range) of simulation time on average (Fig. 2D). We note that these residues have higher occupancies for PIP compared to other lipids, including other negatively charged phospholipids (PA, PS and PI) (Fig. 2D).

Simulations in a complex mammalian membrane showed that PIP species bind specifically and selectively to Na_v_1.4 in the presence of other negatively charged lipids and at low, physiological concentrations. However, the large membrane required, and long PIP binding durations prevented sampling of large numbers of binding and unbinding events. To address this, we carried out additional simulations where Na_v_1.4 was placed in a smaller POPC membrane with a 5% concentration of each PIP species in the cytoplasmic leaflet (Fig. 3A).

**Fig. 3.**
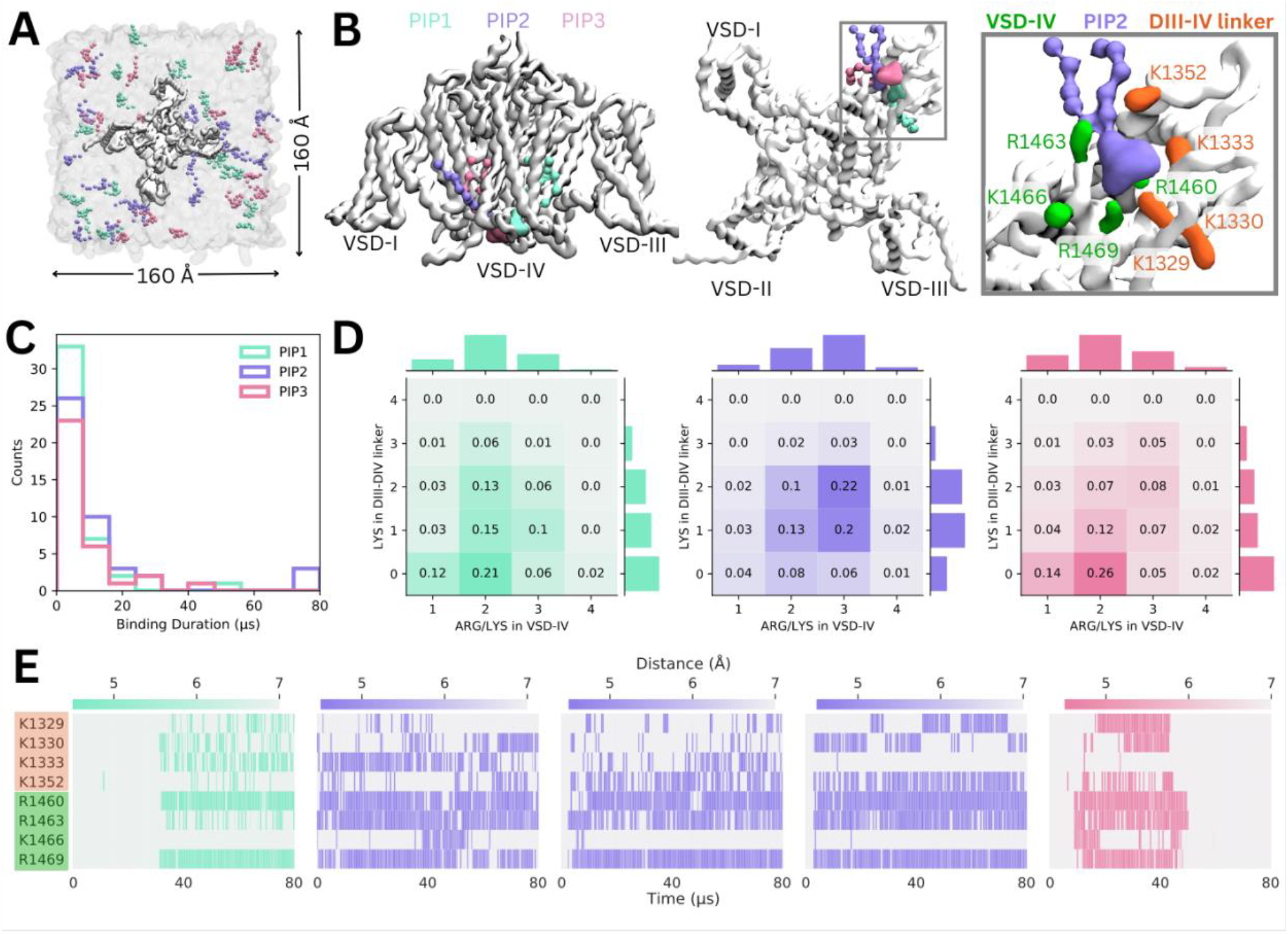
Binding of different PIP species in enriched PIP simulations. **(A)** Enriched PIP simulation system, with Na_v_1.4 embedded in a POPC membrane (transparent gray) and 5% each of PIP1 (blue), PIP2 (purple) and PIP3 (pink) added to the cytoplasmic leaflet. **(B)** Representative snapshots from the five longest binding events from different replicates, showing the three different PIP species (PIP1 in blue, PIP2 in purple and PIP3 in pink) binding to VSD-IV and the DIII-IV linker. Na_v_1.4 is shown in white with interacting residues on the DIV S4-S5 linker and the DIII-IV linker colored in green and orange respectively. **(C)** A frequency distribution showing interaction times for each PIP species, defined as the length of a continuous period in which a PIP was within 0.7 nm of two VSD IV binding site residues. **(D)** Frequency plots showing number of positive residues interacting with bound PIP in the DIII-IV linker (vertical) and VSD-IV (horizontal). **(E)** Minimum distance between binding residues on Na_v_1.4 and bound PIPs lipid across simulation time for the five longest binding events, colored by distance and the type of PIP bound.

Across ten replicates of these enriched PIP simulations, each carried out for 80 μs, we observed all three PIP species binding at the identified site on VSD-IV and DIII-IV linker (Fig. 3B). PIPs can approach and bind to this site from either side of the VSD, however, PIP1 only forms stable interactions when it approaches and binds from the VSD-I side while PIP2 and PIP3 usually bind from the VSD-III side (Fig. 3B). These enriched PIP simulations also revealed additional positively charged residues in the DIII-IV linker (K1329, K1333 and K1352) and DIV S4 (R1460) which support binding.

There were 156 PIP binding/unbinding events with duration greater than 2 μs occurring in the identified site (Fig. 3C). Of these, 43 were with PIP1, 44 with PIP2 and 33 with PIP3. The number of short-term (2-10 μs) interactions decreased with headgroup charge. That is, PIP1 formed the greatest number of short-term interactions while PIP3 had the fewest. Of the 31 binding events with duration greater than 10 μs, 7 were with PIP1, 15 with PIP2 and 9 with PIP3 (Fig. S4-5).

When we analyzed interactions occurring during 2-80+ μs binding events, we found that the number of interacting basic residues changes depending on the PIP headgroup charge (Fig. 3D). PIP1 (headgroup charge: -3e) binding is most frequently coordinated by two positive charges in VSD-IV and zero or one residue in the DIII-IV linker. PIP2 (headgroup charge: -5e) binding most frequently involves one or two interactions from the DIII-IV linker and three from VSD-IV for a total of five interactions. Interestingly, despite its greater negative charge, PIP3 (headgroup charge: -7e) interacts similarly to PIP1, and has fewer interactions than PIP2.

The minimum distance between the interacting PIP headgroup and each binding residue across simulation time is shown for the five 40+ μs PIP binding events (Fig. 3E). Of these, three are PIP2 binding events which almost span the entire 80 μs of simulation. One 40+ μs binding event each for PIP1 and PIP3 were also observed. The stable PIP1 binding event observed involved interactions with R1460 and R1469 in VSD-IV, as well as fluctuating interactions with the four residues of the DIII-IV linker (K1329, K1330, K1333, K1352). The three long term PIP2 binding events observed were similar to each other, mainly stabilized by R1460, R1463 and R1469. As with PIP1, the number and identity of interacting DIII-IV linker residues varied across the span of each simulation and between replicates, owing to the flexibility of the linker. Like PIP2, PIP3 binding was characterized by stable interactions with R1460, R1463 and R1469 (VSD-IV) as well as K1329 and K1330 (DIII-IV linker). *In silico* mutation of the eight residues implicated in PIP binding to leucine (charge neutralization while preserving sidechain size) or glutamate (charge reversal) significantly reduced PIP binding (Fig. S6).

Coarse-grained simulations enhance sampling by reducing the number of particles, enabling larger time steps and construction of a smoother energy surface. To examine interactions and protein conformational changes in atomistic detail, we backmapped representative snapshots from our PIP enriched simulations, where we observed stable, long-term binding events between Na_v_1.4 and PIP1/PIP2. The coarse-grained PIP1 and PIP2 molecules were converted to the atomistic PI(4)P and PI(4,5)P_2_ respectively. For each system, 7.5 μs of atomistic simulations were performed (5 replicates, 1.5 μs each).

In these simulations, the PI(4,5)P_2_ headgroup was stable at the binding site identified in coarse-grain simulations (RMSD <2.2 Å) (Fig. 4A, Fig. S7). PI(4,5)P_2_ binding is predominantly coordinated by R1469 on the DIV S4-S5 linker, as well as R1466 and R1463. In one replicate, PI(4,5)P_2_ associates with R1460 via the phosphate group (PO4) that connects the headgroup to the PIP tails. The PI(4,5)P_2_ headgroup also forms electrostatic interactions with K1329 and K1330 in the DIII-IV linker, but forms few contacts with K1333 (Fig. 4B, Fig. S8). This loop portion of the DIII-IV linker is highly flexible (Fig. S7), thus the lysine sidechains are able to flip between binding the PI(4,5)P_2_ headgroup and facing the intracellular space.

**Fig. 4.**
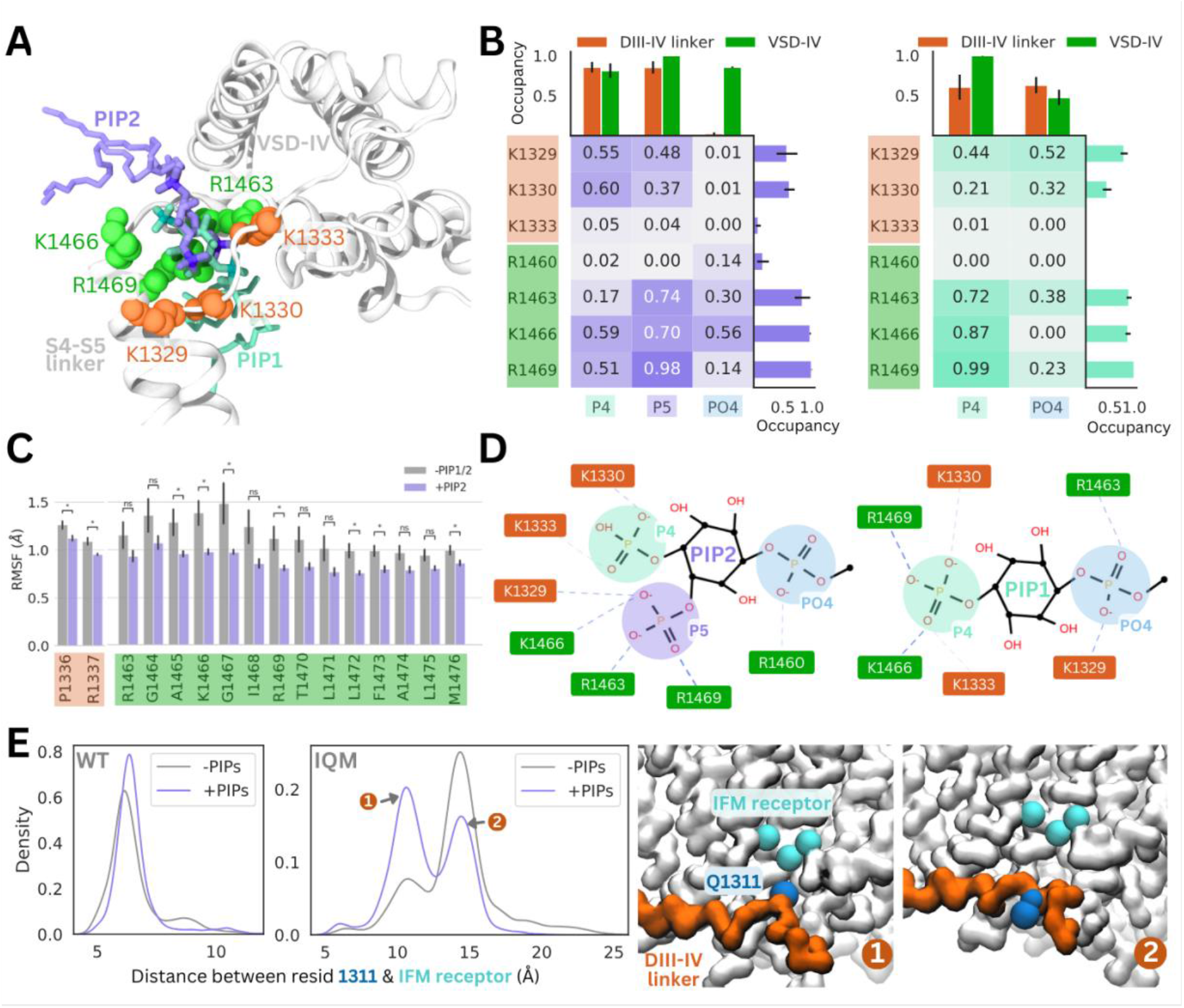
PI(4,5)P_2_ and PI(4)P binding to Na_v_1.4 stabilizes the DIII-IV linker in atomistic and flexible coarse-grained simulations. **(A)** Representative snapshots of PI(4,5)P_2_ bound from the VSD-I side (purple stick) and PI(4)P bound from the VSD-III side (cyan stick), with six basic residues forming the binding site located on the DIII-IV linker (orange VDW representation) and VSD-IV S4-S5 linker (shown in green VDW representation) visualized from the intracellular face of the protein **(B)** Proportion of frames where each of the binding site residues were identified to be within 4.5 Å with the different headgroup regions, P4, P5 and PO4, for PI(4,5)P_2_ (left) and PI(4)P (right) **(C)** Comparison of RMSF per carbon-alpha for simulations with and without bound PI(4,5)P_2_, showing residues on the S4-S5 linker and DIV linker with significant differences in mobility (p-value < 0.05) **(D)** Interaction network plots between the PIP headgroup and basic binding residues on DIII-IV linker (orange) and DIV S4-S5 linker (green), generated by ProLIF – showing the dominant interactions across simulations of PI(4,5)P_2_ and PI(4)P. **(E)** Density plots showing differences in the distributions of distance between IFM/IQM motif and its binding pocket in the presence and absence of PIPs for the Nav1.4 wild-type (left) and IFM->IQM mutant (right); with representative snapshots showing the two distinct conformations of the IQM motif in the mutant.

The 4’-phosphate formed interactions with residues belonging to the DIII-IV linker (K1329 and K1333) and DIV S4-S5 linker (K1666 and R1469) with similar frequency, in 55-60% of simulation frames (Fig. 4B). In contrast, the 5’-phosphate formed contacts with three VSD-IV S4-S5 residues (R1463, K1466 and R1469) in 70-98% of simulation frames and DIII-IV linker residues in 37-48% of frames. Taken together, this data suggests that although the headgroup is flexible when bound, the 5’-phosphate is more important for coordinating VSD-IV S4-S5 residues while the 4’-phosphate associates with both regions of the binding site.

Atomistic simulations of PI(4)P bound from the VSD-III side show that this is also a stable pose (RMSD <2 Å), where the headgroup interacts with the same positively charged residues as seen for PI(4,5)P_2_ (Fig. 4A, Fig. S7). The residues on the S4-S5 linker, R1463, K1466 and R1469, predominantly bind to the 4’-phosphate (Fig. 4B). Due to the more buried location of PI(4)P binding, the PO4 phosphate can associate more with DIII-IV linker lysines (Fig. 4B), however, the absence of the 5’-phosphate leads to a reduced number of total electrostatic interactions (Fig. S9).

To investigate structural changes that might occur in the presence of PI(4)P/PI(4,5)P_2_, we also carried out simulations of the inactivated Na_v_1.4 structure without any PIPs bound for comparison. Although the DIII-IV linker remains bound throughout simulations both with and without PIPs, residues P1336 and R1337 in the DIII-IV linker downstream of the IFM motif are significantly less mobile with PI(4,5)P_2_ present (Fig. 4C). Additionally, seven residues belonging to the DIV S4-S5 linker, including binding residues K1466 and R1469, also have significantly lower mobility when PI(4,5)P_2_ is bound (Fig. 4C). In the PI(4)P bound simulations, there were no significant differences in DIII-IV linker or S4-S5 linker mobility.

To further probe whether PIP can stabilize the DIII-IV linker and the inactivation gate, we applied coarse-grained simulations with the DIII-IV linker unrestrained. In simulations of WT Nav1.4, the IFM has a reduced stability within its binding pocket when PIP is excluded from the membrane (Fig. 4E). To accentuate this effect we simulated an inactivation-deficient variant of Nav1.4, where the IFM motif is mutated to IQM (26). We find that the IQM motif has a greater probability of being tightly associated with the receptor pocket in the presence of PIP compared to without it (Fig. 4E), supporting our observation that PIP can stabilize the channel in the inactivated state. This suggests that the presence of PIP may partially rescue some of the structural defects associated with inactivation dysfunction in Nav mutants.

The PIP binding poses seen in our simulations is similar to resolved binding poses for PI(4,5)P_2_ in cryo-EM structures of Ca_v_2.2 and K_v_7.1 (Fig. 5A). A sequence alignment of Na_v_1.4 VSDs show that there are more positively charged residues present in the S4/S4-S5 linker regions of DIV compared to the other domains (Fig. 5B). PIP is bound at similar VSD residues on both these ion channels, with PIP forming interactions with a gating charge further up the S4 helix in Ca_v_2.2 due to the VSD being in a different state. Additionally, the high sequence similarity in the S4/S5-S5 linker and DIII-IV linker regions between the nine human Na_v_ channel subtypes suggests conservation of the PIP binding site (Fig. 5C).

**Fig. 5.**
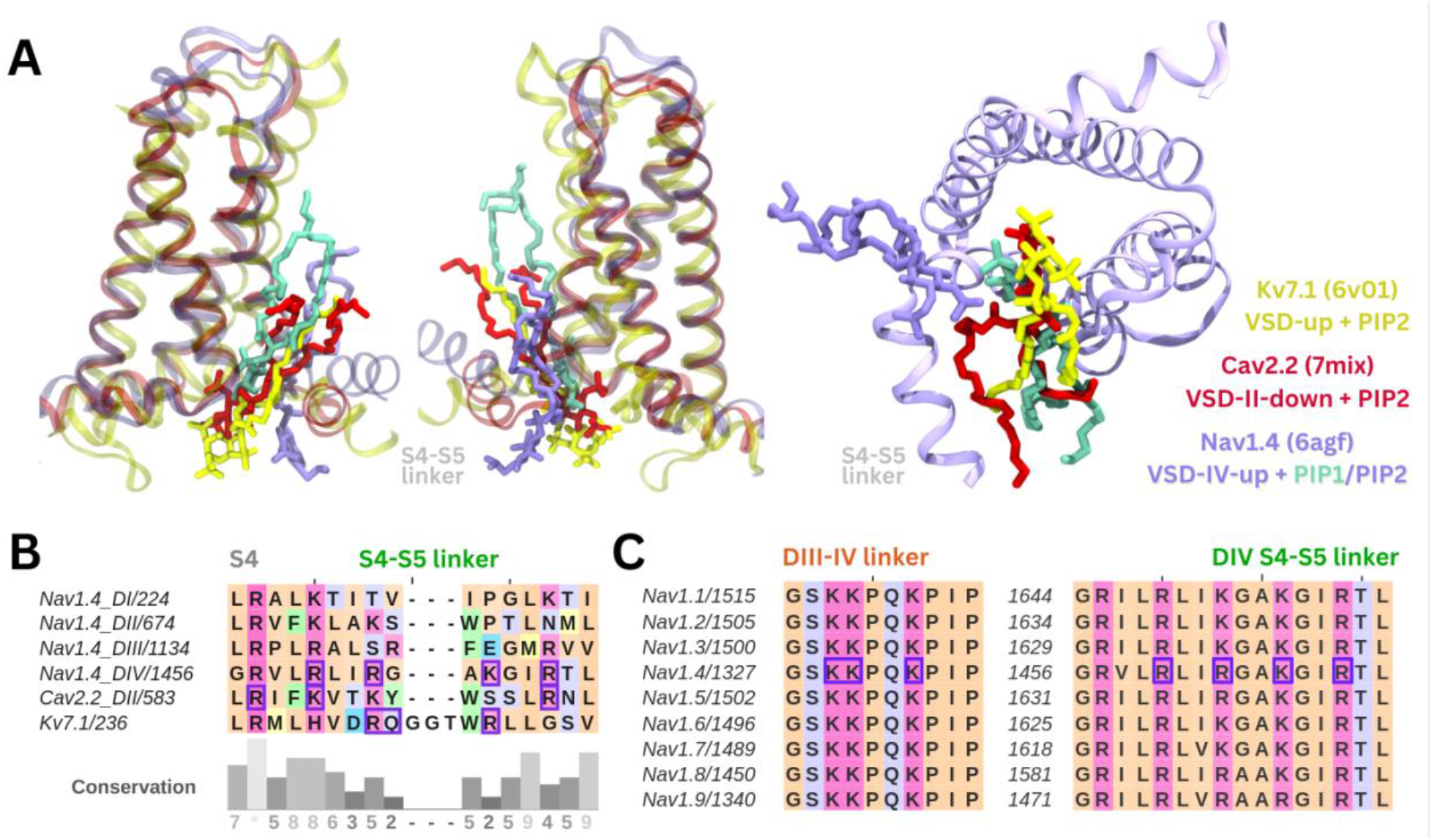
Comparison of the identified phosphoinositide binding site to Na_v_ subtypes and other ion channels. **(A)** Binding poses of PI(4,5)P_2_ (in purple) and PI(4)P (in cyan) aligned with two other tetrameric channels structures K_v_7.1 (6v01, in yellow) and Ca_v_2.2 (7mix, in red) that were resolved with PI(4,5)P_2_ at their respective VSDs. **(B)** Sequence alignment of the S4 helix and S4-S5 linker of the four domains of Na_v_1.4, compared to VSD-II of Ca_v_2.2 and one of the four identical VSDs of K_v_7.1; residues colored by amino acid class; purple boxes indicate PI(4,5)P_2_ binding residues (identified with 5 Å of the headgroup). **(C)** Sequence alignment of the nine human Na_v_ channel subtypes shows high sequence similarity in the S4 helix, S4-S5 linker and DIII-IV linker regions.

To assess possible state- and subtype-dependent differences in PIP binding, we simulated three structures of Na_v_1.7 with different VSD conformations in a PIP enriched membrane (Fig. 6A). The inactivated Na_v_1.7 structure (blue, PDB ID: 6j8g) contains all VSDs in the activated/up state and a bound DIII-IV linker. We also simulated a Na_v_Pas (American cockroach) chimera structure with a human Na_v_1.7 VSD-IV in the deactivated/down state (VSDs I-III are activated, and from Na_v_Pas) (pink, PDB ID: 6nt4). This structure (14) features a dissociated DIII-IV linker and a resolved C-terminal domain bound to the DIV S4-S5 linker (forming the ‘inactivation switch’) at residues identified to form part of our PIP binding site. To assess PIP binding at VSD I-III in the deactivated/down state, we modelled the Na_v_1.7 resting state based on templates structures in which different VSDs have been captured in deactivated states with the aid of toxins (detailed in Table S2). These structures feature three or more of the gating charge residues below the hydrophobic constriction site (HCS) and displacement of the S4-S5 linker (Fig. S10). Given that no resting state mammalian Na_v_ channel structure has been resolved, it is possible that the modelled VSDs may not reflect the fully deactivated state.

**Fig. 6.**
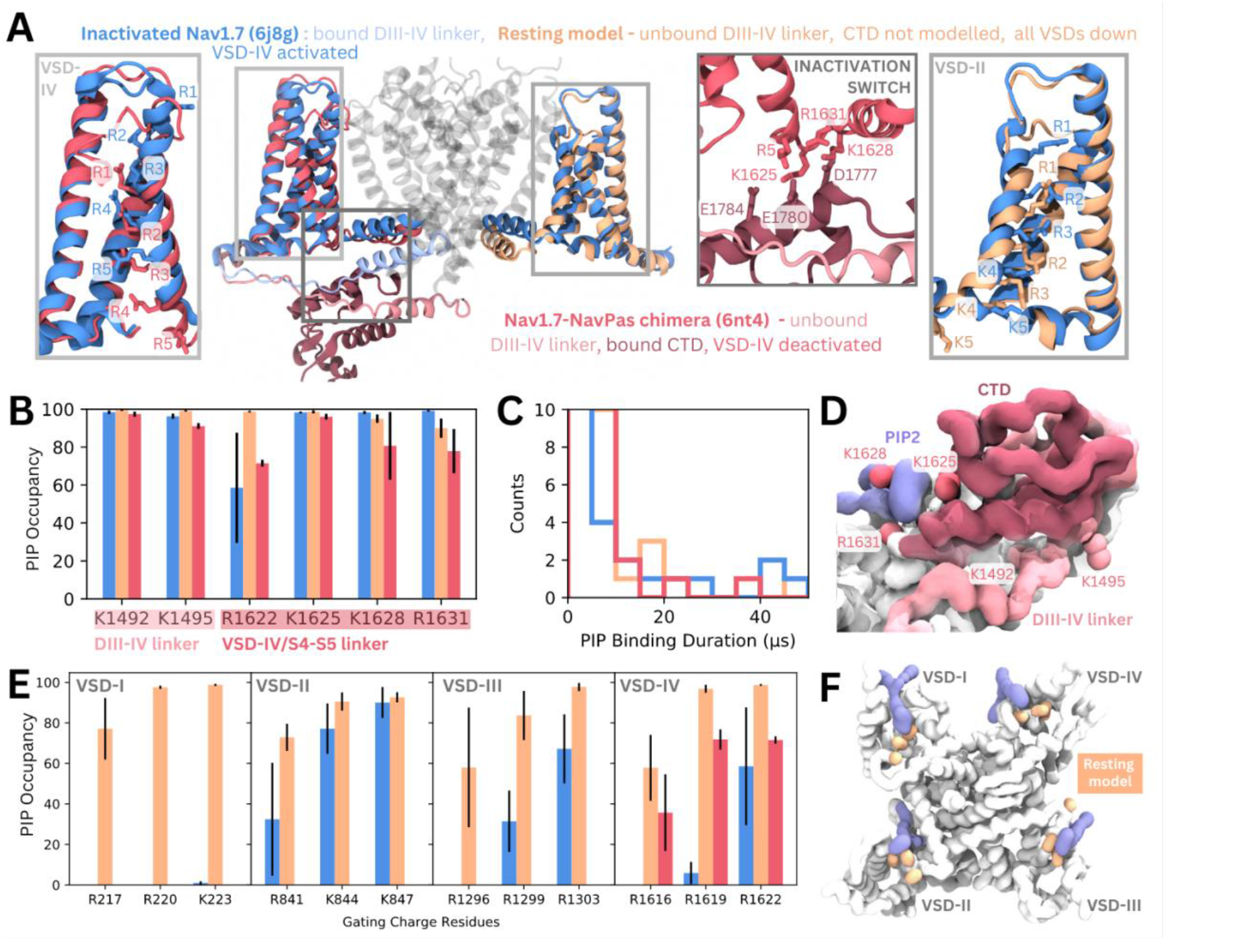
PIP binding to Na_v_1.7 with different VSD states in coarse-grained simulations. **(A)** Atomistic representation of the three different Na_v_1.7 structures simulated: (1) the inactivated state (blue, PDB ID: 6j8g) with the VSDs all in the activated, up state (2) the Na_v_1.7-Na_v_Pas chimera (pink, PDB ID: 6nt4) with the Na_v_1.7 VSD-IV in the deactivated, down state and a bound Na_v_Pas CTD (3) a Na_v_1.7 resting state model (orange, model generation detailed in Table. S2) with all four VSDs in the deactivated, down state. Panel insets show the different conformations of VSD-IV (left) and VSD-II (right) across different structures. The inactivation switch formed by the CTD and VSD-IV S4-S5 linker proposed by Clairfeuille et al. is shown (middle). **(B)** Combined occupancy of all PIP species (PIP1, PIP2, PIP3) at binding site residues in the three systems **(C)** Distribution of PIP binding durations at the identified site **(D)** Intracellular view of CTD covering the resting state VSD-IV. Representative snapshot of PIP binding at DIV S4-S5 linker in the Na_v_1.7-Na_v_Pas system. The CTD (dark pink) prevents PIP access to DIII-IV linker lysines, K1492 and K1495. **(E)** Combined PIP occupancy at the bottom three gating charges on VSD I-III in the inactivated (blue) and resting state model (orange) simulations. For VSD-IV, PIP occupancy in the Na_v_1.7-Na_v_Pas system (pink) is also shown. **(F)** Representative simulation snapshot showing PIP (purple) binding at the gating charges (orange) in the resting state model simulations.

In triplicate 50 μs coarse-grained simulations, PIPs bind to the analogous site to that seen in inactivated Na_v_1.4 in the inactivated Na_v_1.7 structure, interacting with residues belonging to both the DIII-IV linker and VSD-IV for durations comparable to Na_v_1.4, (Fig. 6B-C, Fig. S11). Binding of PIP to the DIV S4-S5 linker to the deactivated VSD-IV in the Na_v_1.7-Na_v_Pas chimera and resting state model was also observed (Fig. 6B-C). However, in the Na_v_1.7-Na_v_Pas chimera, the PIP bound at the S4-S5 linker cannot simultaneously associate with the DIII-IV linker (Fig. S11), due to its sequestration by the C-terminal domain, which moves the lysines away from the binding VSD-IV residues (Fig. 6D). Instead, K1491, K1492 and K1495 (on the DIII-IV linker) are occupied by different PIPs on the other side of the VSD. In the resting state model, which features an unbound DIII-IV linker PIPs binding at the DIV S4-S5 residues also do not associate with the DIII-IV linker (Fig. S11). Comparison of binding durations at this site across the three systems reveals a greater number of long PIP interactions (> 20 μs) with inactivated Na_v_1.7 (5) compared to Na_v_1.7-Na_v_Pas (3) or the resting state model (2) where VSD-IV is down, and the DIII-IV linker is dissociated.

There are state-dependent differences in PIP occupancy at the gating charges in each VSD (Fig. 6E-F, Fig. S12A-B). PIP can associate with the lowest three gating charges VSD-I of the resting state model, but not in the inactivated state (Fig. 6E). This is likely due to the large displacement in the DI-S4 helix, which moves down three helical turns in the resting state model so that all gating charges are below the HCS (Fig. S10). In VSD II and III, the PIP interaction differences between inactivated and resting state are present but less pronounced, owing to a smaller difference in the relative displacement between the gating charges between states (Fig. S10). PIP occupancy is also higher at VSD-IV when it is in the deactivated conformation however, the presence of a bound CTD as seen in the Na_v_1.7-Na_v_Pas model, reduces this occupancy of PIP (Figure 6E). Additional simulations of a resting state model of Nav1.4 built using our Nav1.7 resting state model as a template suggest that similar gating charge interactions occur for Nav1.4 when the VSDs are deactivated (Fig. S12C).

## Discussion

Recently, PI(4,5)P_2_ was shown to be a negative regulator of Na_v_1.4, modulating channel kinetics and voltage dependence. Presence of PI(4,5)P_2_ causes a depolarizing shift in the voltage dependence of activation, that is a stronger stimulus is required to produce Na_v_1.4 opening. Additionally, it stabilizes the inactivated state of Na_v_1.4, marked by both shortened times to inactivation and slowed recovery from the inactivation.

Using a multiscale simulation approach, we identified a putative PIP binding site comprised of positively charged residues belonging to the S4 helix/S4-S5 linker of VSD-IV (R1460, R1463, K1466, R1469) and DIII-IV linker (K1329, K1330, K1333, K1352). Coarse grained simulations of Na_v_1.4 embedded in a complex membrane showed that PIP interacts with residues belonging to VSD-IV and the DIII-IV linker. In coarse grained enriched PIP simulations, PIP2 formed longer duration interactions with Na_v_1.4 than PIP1 and PIP3, supported by a greater number of charged interactions. Atomistic simulations verified the stability of PI(4,5)P_2_ (the most common PIP2 species in the plasma membrane) at this site and showed that the binding of PI(4,5)P_2_ reduces the mobility of some DIV S4-S5 and DIII-IV linker residues. Simulations of Na_v_1.7 with VSDs in different conformational states showed that the PIP binding site is conserved in Na_v_1.7 and that PIP interactions at VSD gating charges are functional state dependent, with more interactions being formed when the VSDs are deactivated.

The DIII-IV linker, CTD and S4-S5 linkers all play key roles throughout the Na_v_ conformational cycle. Mutation of the IFM inactivation motif as well as other residues in the DIII-IV linker alter fast inactivation and recovery from fast inactivation (27, 28). While the precise role of the Na_v_ C-terminal domain and its conformation during the Na_v_ activation cycle remain elusive, it is likely to be important for coordinating fast inactivation (14, 29). CTD binding to DIV S4-S5 and sequestration of the DIII-IV linker is proposed to occur in the resting state. After the pore opens, activation of VSD-IV is thought to cause CTD dissociation, releasing the DIII-IV linker to allow fast inactivation. Residues in ‘switch 1’ of the CTD-binding site on the DIV S4-S5 stably bind PIP in our simulations of inactivated Na_v_1.4 and Na_v_1.7. We hypothesize that PIP binding at this location makes it more difficult for the CTD to reassociate with VSD-IV, a conformational change which is required during recovery from inactivation.

More generally, the S4-S5 linkers in all four domains couple the VSD to the pore helices and adopt different orientations depending on VSD activation states. When the VSD is activated (in the open and inactivated states), the S4-S5 linkers lie parallel to the membrane. In the resting state, when the voltage sensor is deactivated, the S4-S5 linkers move downward below the plane of the membrane. We propose that PIP binding at the identified site could additionally stabilize both the DIV S4-S5 linker and DIII-IV linker to favor the inactivated state. Although the recovery from fast inactivation occurs on the order of several milliseconds (30), and is beyond atomistic simulation timescales, we observed statistically significant reductions in the RMSF of several DIII-IV linker and DIV S4-S5 linker residues when PI(4,5)P_2_ was bound in 1.5 μs atomistic simulations. Reduction in the mobility of the DIII-IV linker may slow the dissociation of the upstream IFM motif and stabilization of the S4-S5 linker prevents the downward movement required for the channel to transition back to the resting state.

The PIP binding residues identified here are conserved in Na_v_1.1-1.9 (Fig. 5C), suggestive of a shared binding site and mechanism for PIP-mediated modulation across subtypes. Mutations at these conserved residues in other subtypes lead to various gain-of-function diseases (Table S1). For example, analogous to the Na_v_1.4 R1469 residue, the R1642C mutation (in Na_v_1.3) leads to developmental epileptic encepahlopathy (31), and R1644C/H mutations in Na_v_1.5 (analogous to R1469 in Na_v_1.4) cause cardiac arrythmias, characterized by accelerated rates of channel recovery from inactivation (32). Consistent with our observations, these diseases with mutations on the DIII-IV linker are likely to reduce PI(4,5)P_2_ binding, which could be a contributing factor to instability of inactivated state in these pathogenic variants. These inactivation-deficient variants, as well as the IQM variant that we simulated, further emphasize that interactions between PI(4,5)P_2_ with Na_v_ channels could be important for prolonging the fast-inactivated state.

The PIP binding site identified here harbors sequence and structural similarity to PI(4,5)P_2_ binding sites found in other cation channels (Fig. 5B). For example, PI(4,5)P_2_ is resolved at a similar site near the VSD and S4-S5 linker in a recent cryo-EM structure of K_v_7.1, where the phosphate headgroup forms analogous contacts to R249 and R243 (PDB ID: 6v01) (20). Despite differences in the role of PI(4,5)P_2_, which negatively regulates Na_v_1.4 but is required for K_v_7.1 pore opening, the binding site appears to be conserved. Based on the PI(4,5)P_2_ binding site, a structurally similar compound was developed as an activator of K_v_7 channels and proposed to be a future antiarrhythmic therapy (33).

PI(4,5)P_2_ also binds to the down, deactivated state of VSD-II in Ca_v_2.2 (PDB ID: 7mix) (21). In this structure, the PI(4,5)P_2_ headgroup interacts with two VSD-II gating charges, R584 and K587. Compared to the positioning of PI(4,5)P_2_ in our simulations of Na_v_1.4 with an activated VSD-IV, the headgroup associates further up the S4 helix in Ca_v_2.2 due to the VSD being in a deactivated state. This is also seen in K_v_7.1 which contains an extended GGT loop in the S4-S5 linker which prevents PI(4,5)P_2_ binding in the VSD-down state (34). In our coarse-grained simulations of the resting Na_v_1.7 model, we observe a similar state-dependent difference in PI(4,5)P_2_ interactions with the deactivated states of each VSD. Since activation of VSDs I-III are known to be coupled to channel opening (17), we propose that PIP binding at these VSDs impedes their ability to activate and thus increases the voltage threshold required of opening. PI(4,5)P_2_binding at VSD-IV is prevented by presence of the CTD (and absence of the DIII-IV linker) in the resting state, thus not affecting the kinetics of inactivation onset.

The leftward shift in voltage dependence of inactivation is less pronounced when PI(4,5)P_2_ is converted to PI(4)P rather than completely dephosphorylated to PI (25). This suggests that PI(4)P may play a compensatory role when PI(4,5)P_2_ is not present. This is supported by our simulations which show that PI(4)P can also stably occupy the identified binding site, albeit with shorter duration and form less electrostatic interactions compared to PI(4,5)P_2_. Our atomistic simulations also showed that the 5’-phosphate is more important than the 4’-phosphate for forming interactions with the DIV S4-S5 linker residues of the inactivation switch. These factors suggest that PI(4,5)P_2_ binding is preferred over PI(4)P at this site and can better compete to bind over the CTD, implying that PI(4,5)P_2_ is more effective at stabilizing the inactivated state and inhibiting recovery to the resting state.

Simulations using the Martini2.2 forcefield have previously been used to investigate lipid-protein interactions (35) and successfully predict specific lipid binding sites, including the PIP binding site on K_ir_ channels (36). While the Martini2.2 PIP species can be parameterized for a specific sub-species (e.g. PI(3,4)P_2_ vs PI(4,5)P_2_), we instead employed atomistic simulations to complement and strengthen findings from coarse grain simulations, allowing us to identify the specific contribution of the 4’- and 5’-phosphate groups to binding as well as investigate the conformational changes associated with PI(4,5)P_2_ binding. Since the precise protonation of the PI(4,5)P_2_ headgroup in a physiological setting is unclear, we explore one case with the 4’-phosphate protonated. Given that the PI(4,5)P_2_ headgroup can adopt slightly different binding orientations and can fluctuate over the course of atomistic simulations (Fig S8), we expect the alternate protonation state to have similar affinity for the binding site.

In this work we made use of the Martini2.2 model for our coarse-grained simulations, however recently the refined Martini3 (37) has become available and will be a useful tool for further interrogating protein-lipid interactions, as a greater number of lipid parameters become available. Our coarse-grained simulations also allow us to investigate the association of other lipid types with Nav1.4. While we focus on PIP here, there are other lipid species that have modulatory effects on Nav channels, such as cholesterol (38), glycolipids, DG, LPC and PI, and their interactions warrant further investigation.

## Conclusion

Using multiscale simulations, we show that PI(4,5)P_2_ binds stably to inactivated structures of Na_v_1.4 and Na_v_1.7 at a conserved site within the DIV S4-S5 linker. As the CTD is proposed to also bind here during recovery from inactivation, we hypothesize that PI(4,5)P_2_ competes with the CTD to bind to this site, prolonging inactivation. At this site, PI(4,5)P_2_ simultaneously binds to the DIII-IV linker which is responsible for allosterically blocking the pore during fast inactivation (Fig. 7). Its binding reduces the mobility of both the DIV S4-S5 and DIII-IV linkers, potentially slowing the conformational changes required for the channel to recover to the resting state. We also propose that in the resting state, PIPs form additional interactions with S4 gating charges, particularly in VSD-1, anchoring them to the membrane in a way which may make the upwards movement required for their activation more difficult. Our results provide insight into how sodium channels are modulated by phosphoinositides, an important step for the development of novel therapies to treat Na_v_-related diseases.

**Fig. 7.**
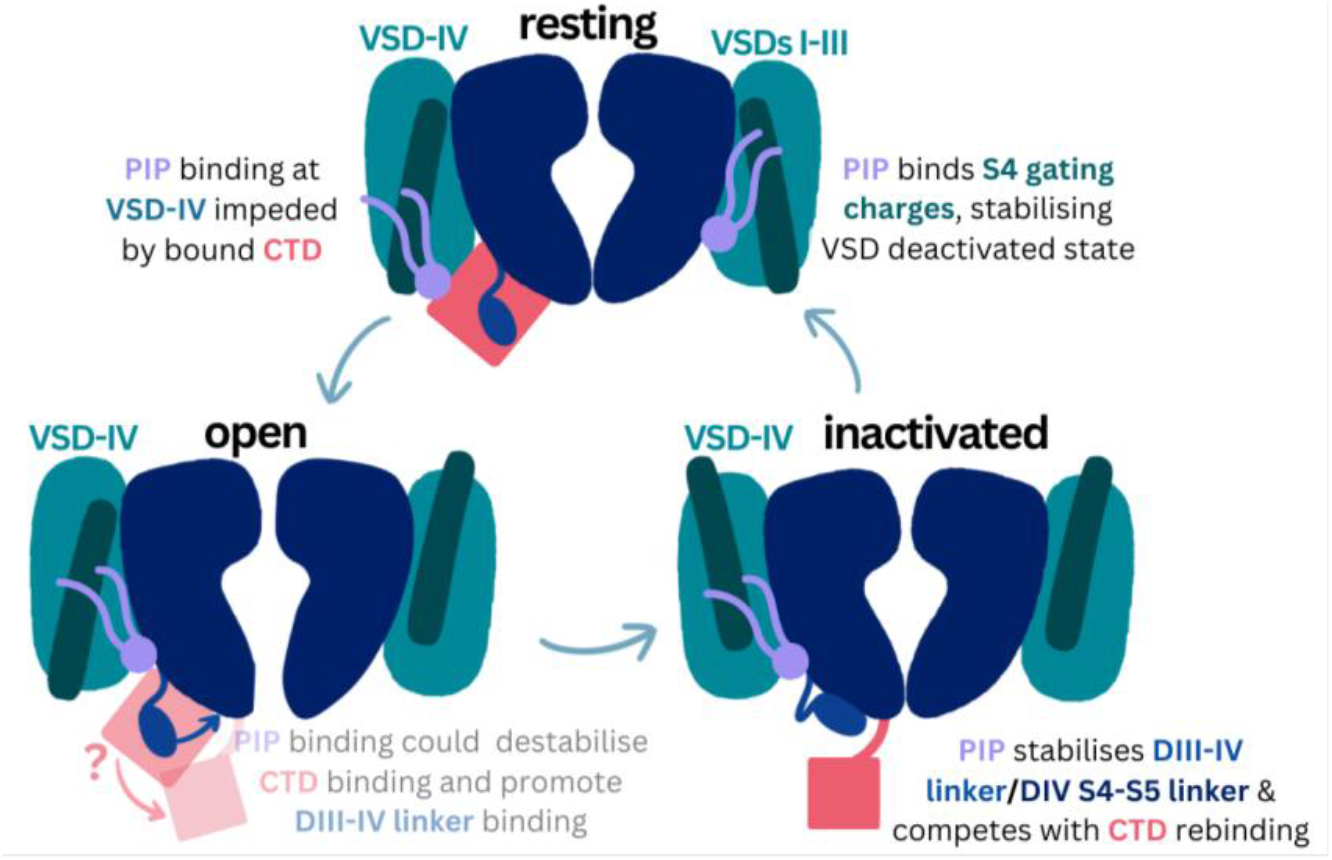
Proposed mechanism of PIP effects on the sodium channel functional cycle.

## Methods

### Coarse grained simulations

Coarse-grained simulations of Na_v_1.4 embedded in a complex mammalian membrane were carried out to investigate lipid-protein interactions. The inactivated Na_v_1.4 alpha subunit (PDB ID: 6agf) (8) was coarse-grained using the CHARMM-GUI Martini Maker (39, 40) and embedded in a 360 Å x 360 Å complex membrane using insane.py (41). The composition of the complex mammalian membrane is as reported in Ingólfsson, *et al*. (42). Three replicate simulations, each with different starting coordinates, were carried out for 16 μs each.

To better sample binding events, we also carried out PIP-enriched simulations in which Na_v_1.4 was embedded in a 160 Å x 160 Å POPC membrane with 5% of each PIP species, PIP1, PIP2, PIP3 (with charge parameters of -3e, -5e and -7e respectively), added to the cytoplasmic leaflet using insane.py. Ten replicate simulations, each with different starting coordinates, were carried out for 80 μs each. To validate our proposed binding site, we additionally mutated the positively charged PIP binding site residues K1329, K1330, K1330, K1352, R1460, R1463, K1466 and R1469 to leucines (‘8L’) or glutamates (‘8E) and simulated these mutant channels in enriched PIP membranes for 20 μs in triplicate. To explore the possibility for PIP to stabilize the DIII-IV linker in an inactivation-deficient Nav1.4 variant, additional coarse-grained simulations were carried out where the DIII-IV linker region (residues L1305-K1341) was unrestrained for both the WT Nav1.4 and simulations in which the IFM motif was mutated to IQM. In these simulations, an elastic network was applied to E1314-G1327 in the linker to preserve the helicity of this region. Flexible linker simulations were conducted in triplicate for 20 μs in both PIP-enriched bilayers and POPC-only bilayers.

To explore possible state and subtype dependent differences in PIP binding, the inactivated Na_v_1.7 structure (PDB ID: 6j8g) (43) and the Na_v_1.7-Na_v_Pas chimera with CTD bound and VSD-IV in the deactivated state (PDB ID: 6nt4) (14) were also coarse-grained and simulated in PIP-enriched membranes (same protocol as above) for three replicates of 50 μs each. Additionally, a model Na_v_1.7 with all four VSDs in the deactivated state was built using Modeller (44) and simulated in triplicate for 50 μs each. The template and structural information for this model are detailed in Table S2. In brief, VSD-I down was modelled from 7xve (45), VSD-II and VSD-III were both modelled from the deactivated VSD-II from 7k48 (46), and VSD-IV and the unbound DIII-IV linker were modelled from the corresponding regions in 6nt4 (14). Adjacent S5/S6 regions to each VSD were also modelled from each specified template to ensure proper contacts between the pore domain and VSDs. The CTD was not included in the model. Using these Na_v_1.7 templates, the resting state model of Na_v_1.4 was generated and simulated for three replicates in coarse grain with the protein backbone restrained, in a PIP-enriched membrane for 50 μs.

All systems were solvated and ionized with 150 mM NaCl. All coarse-grained simulations were carried out with GROMACS 2022 (47) using the Martini2.2 forcefield (48) and the PIP parameters for each charge state, where PIP1 is based on PI(3)P and PIP2 is based on PI(3,4)P2 (49). Energy minimization was carried out on each system using the steepest descent method for 1,000 steps. Following this, equilibration in the constant pressure, constant volume (NVT) ensemble at 1 atm for 10 ps was carried out with backbone position restraints (1,000 kJ mol^−1^nm^−2^) using a 2-fs timestep. Following this, constant pressure and temperature (NPT) equilibration simulations were carried out, using 5-, 10-, and 20-fs timesteps in sequence, with each running for 5,000 steps. 1 atm pressure was maintained using a Berendsen barostat with semi-isotropic conditions. Production simulations were carried out in the NPT ensemble, kept at a temperature of 310 K using the Nose-Hoover thermostat (50) and a pressure of 1 bar using the Parrinello-Rahman barostat (51). A time step of 20 fs was used. During production simulations, the backbone beads were weakly restrained to their starting coordinates using a force constant of 10 kJ mol^−1^nm^−2^.

### Atomistic simulations

Atomistic simulations were performed to characterize atomic interactions between Na_v_1.4 residues and the bound PI(4,5)P_2_ headgroup. Frames from a stable PIP2 binding event (from replicate 1 of enriched PIP simulations) were clustered using a selection of the bound PIP2 headgroup beads (C1 C2 C3 PO4 P1 P2) and binding residues K1329, K1330, K1333, K1463, K1466 and R1469 with an RMSD cutoff of 2.5 Å. The protein and bound PIP2 were extracted from the representative frame of the cluster and backbone beads of the coarse-grained VSD-IV were aligned to the corresponding carbon-alpha atoms in the original cryo-EM structure of Na_v_1.4 (8). PIP2 was backmapped to atomistic coordinates of SAPI24 (the CHARMM lipid for PI(4,5)P_2_ with -2e charge on P5, -1e charge on P4 and -1e on the PO4, as shown in Fig 4D) and the protein was replaced with the 6agf structure. The system was embedded in a 140 Å x 140 Å POPC membrane, solvated and 0.15 M NaCl added using the CHARMM-GUI Membrane Builder (39, 52, 53). An identical system was set up with Na_v_1.4 in a POPC membrane without PIP.

Atomistic simulations were performed with Amber20 (54), using the CHARMM36m (55) and TIP3P water (56) forcefields. Equilibration steps were performed (minimization, heating, pressurizing), with 5 kJ/mol restraints on the protein backbone, followed by 24 ns of gradually reducing restraints. Five replicates of unrestrained production equilibrium simulations were performed, run for 1.5 μs each. The temperature was set at 310 K using the Langevin thermostat (57) and at a collision frequency of 5 ps^-1^. Pressure was set at 1 bar using the Monte Carlo barostat (58) with anisotropic scaling and relaxation time of 1 ps. 12 Å van der Waals cut-off and hydrogen bond SHAKE constraints were used. Hydrogen mass repartitioning was used to enable a 4 fs timestep (59). PI(4)P_1_ was also simulated with the same atomistic procedures, using the SAPI14 CHARMM lipid (with -2e charge on P4 and -1e on the PO4, as shown in Fig 4D).

### Analysis

Coarse grain lipid-protein interactions were characterized using in-house python scripts which used the numpy, MDAnalysis (60) and pandas libraries (as done previously in Lin, Buyan and Corry (61)). A cut-off of 0.7 nm was used to define interactions between lipids and protein residues. Distance heatmaps were generated based on the minimum distance between a PIP bead and the side chain (SC2) bead of the arginine or lysine residue interacting. Binding durations were calculated by first counting the number of interactions between each PIP and VSD-IV binding residues (R1460, R1463, K1466 and R1469). A binding event is defined as the time between the first and last time that PIP interacts with two or more binding residues on VSD-IV, if it interacts with a minimum of one binding residue between this period. Stability of IFM/IQM motif binding in flexible linker coarse-grained simulations was assessed by measuring the distance between the center of mass of the phenylalanine/glutamine residue to the center of mass of three residues (L1153, I1485 and N1591) within the IFM receptor site.

For atomistic simulations, MDAnalysis (60) was used to calculate RMSD of various parts of the protein and PIP headgroup, and RMSF of the carbon-alphas. Statistical significance in RMSF was assessed using student’s t-test. ProLIF (62) was used to compute electrostatic interactions between the protein and binding residues. Representative snapshot of PIP2 binding was generated using the WMC Clustering Tool in VMD to identify the top cluster of the PIP2 headgroup (RMSD cutoff of 3 Å), then subsequently cluster the six binding residues to identify the most representative binding configuration. Trajectories were strided every 1 ns and the first 250 ns of simulations was discarded as equilibration time for analyses of RMSF, ProLIF interactions and clustering. All analysis scripts are available on <u>GitHub</u>.

Simulations were visualized and protein image figures produced using Visual Molecular Dynamics (VMD) (63). ClustalOmega (64, 65) and JalView (66) were used to generate and visualize sequence alignments. Structural representations of K_v_7.1, Ca_v_2.2 and Na_v_1.4 structures were created in VMD by aligning each of the S4 helices on the VSD where the PIP was bound.

## Acknowledgements

The authors thank Ruitao Jin for his contributions to the resting state homology model generation and comments on the manuscript draft. This research was undertaken with the assistance of resources and services from the National Computational Infrastructure (NCI), which is supported by the Australian Government. This research is also supported by Australian Government Research Training Program (RTP) Scholarships. Y.L. is supported by a 2023 NCI HPC-AI Talent Program Scholarship.

