## Supplementary Information for "A binding site for phosphoinositides described by multiscale simulations explains their modulation of voltage gated sodium channels"

**This PDF file includes:**

**Table S1 – Disease causing mutations at analogous PIP binding residues in $Na_v$ subtypes.**

**Table S2 – Generation of the resting state model using a combination of multiple templates for different domains of $Na_v1.7$.**

**Fig. S1-2 – Lipid Z-Density Maps (left) and Contact Occupancy Structures (right) for all 12 lipid types in the mammalian membrane.**

**Fig. S3 – Contact occupancy distributions (left) and outlying residues (right) for all 12 lipid types in the mammalian membrane.**

**Fig. S4-5 – Minimum distance between binding residues on $Na_v1.4$ and bound PIPs lipid across 31 long duration binding events (>10 μs), colored by distance and the type of PIP bound.**

**Fig. S6 – PIP occupancy at binding site residues when mutated to leucine (charge neutralization) and glutamate (charge reversal).**

**Fig. S7 – Root Mean Square Deviation (RMSD) of the $Na_v1.4$ backbone, VSD-IV backbone, DIII-IV linker backbone and S4-S5 backbone (and PIP2/PIP1 headgroup) over 1.5 μs of atomistic simulations with and without PIP.**

**Fig. S8 – Electrostatic interactions between the PIP2 headgroup and binding residues.**

**Fig. S9 – Electrostatic interactions between the PIP1 headgroup and binding residues.**

**Fig. S10 – $Na_v1.7$ resting state model validation.**

**Fig. S11 – Minimum distance between binding residues on $Na_v1.7$ and bound PIPs.**

**Fig. S12 – PIP binding at S4 gating charges in Nav1.7 and Nav1.4.**

**Table S1 – Disease causing point mutations at analogous PIP binding residues in $Na_v$ subtypes (described in the UniProt database).**

| $Na_v$1.4 RESIDUE # | ANALOGOUS RESIDUE # | SUBTYPE | DISEASE INFORMATION; MECHANISM |
|---|---|---|---|
| K1330 | K1505N | $Na_v$1.5 | Long QT3 syndrome; unknown significance |
| R1463 | K1641N | $Na_v$1.2 | Benign familial infantile seizure; unknown significance |
| R1469 | R1657C | $Na_v$1.1 | Generalised epilepsy with febrile seizures plus; depolarising shift in voltage dependence of activation, reduced current, accelerated recovery from slow inactivation |
| | R1642C | $Na_v$1.3 | Developmental epileptic encephalopathy; accelerated recovery from inactivation |
| | R1644C<br>R1644H | $Na_v$1.5 | Long QT3 syndrome<br>Brugadas syndrome |

**Table S2 – Generation of the resting state model using a combination of multiple templates for different voltage sensor domains (VSDs) and pore domains (PDs) of $Na_v$1.7.**

| Template pdb | Domains Used | Res IDs (from template) | Res IDs (human $Na_v$1.7 numbering) | Template Info |
|---|---|---|---|---|
| 7xve | VSDI, PDI, PDII | 1-404, 541-650 | 8-411, 864-972 | $Na_v$1.7 Mutant (L866F, T870M, and A874F on S5II; V947F, M952F, and V953F on S6II; and V1438I, V1439F, and G1454C on S6III, E156K on S2I and G779R on S2II)<br>VSDI deactivated (VSDII partially deactivated but not used) |
| 7k48 | VSDII, PDII, VSDIII, PDIII, PDIV | 405, 938, 1115-1244 | 728-973, 1175-1462, 1639-1768 | $Na_v$1.7 /$Na_v$Ab chimera, where top half of each VSD is $Na_v$1.7 VSDII<br>All VSDs deactivated with engineered tarantula toxin m3-Huwentoxin-IV bound |
| 6nt4 | VSDIV, PDIV, PDI | 939-1244, 239-404 | 1463-1768, 246-411 | $Na_v$1.7/$Na_v$Pas chimera, full $Na_v$1.7 VSDI and VSDIV deactivated with α-scorpion neurotoxin AaH2 bound |
| 7w9k | Loops between S5 and S6 regions for PDII, PDIII and PDIV | 563-619, 811-918, 1141-1213 | 886-942, 1335-1442, 1665-1737 | Fast-inactivated $Na_v$1.7 (All VSDs activated)<br>DIII-IV linker bound to pore |

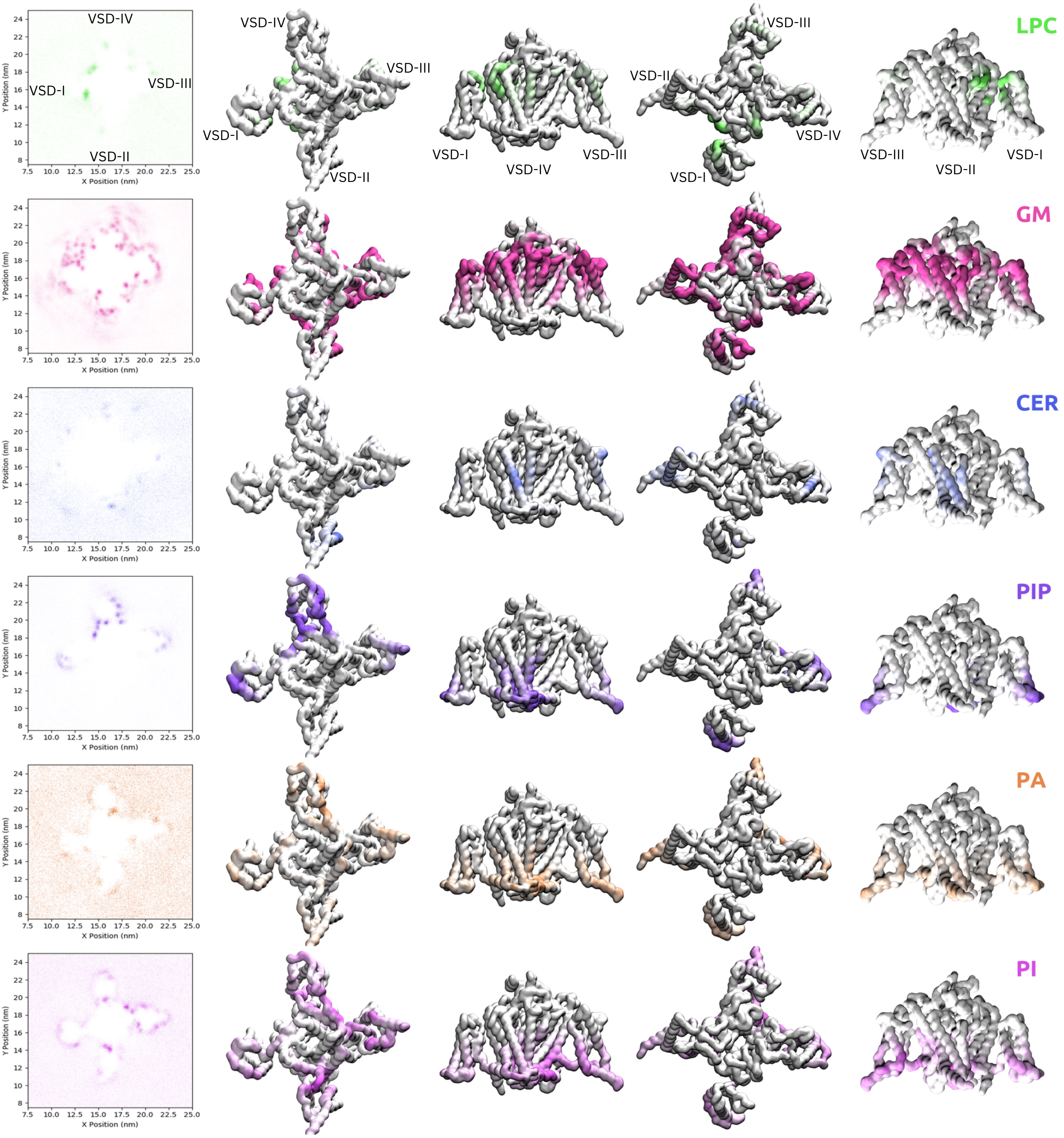

VSD-IV
VSD-I
VSD-III
VSD-II
LPC
GM
CER
PIP
PA
PI
Y Position (nm)
X Position (nm)

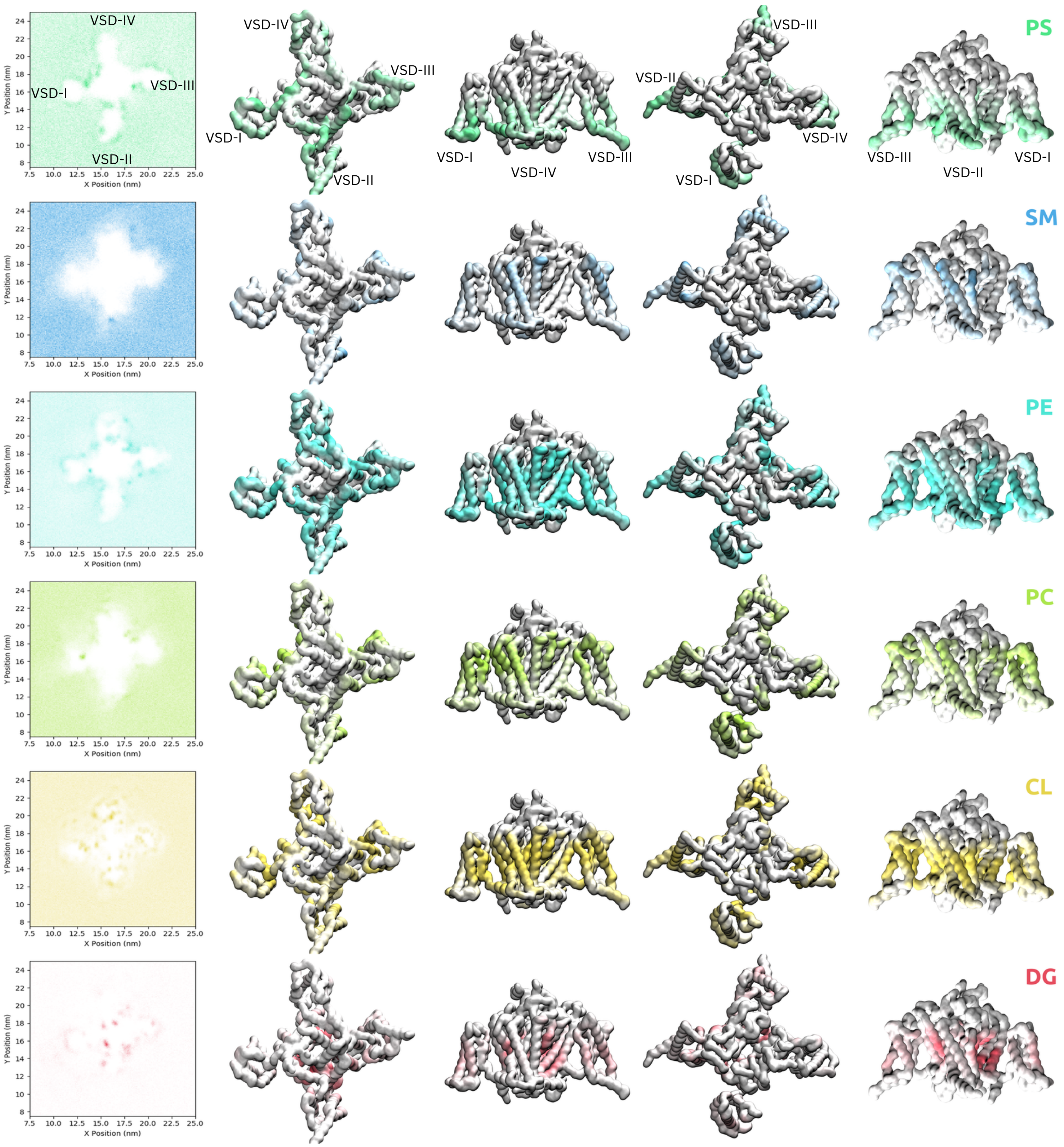


**Fig. S1-2 – Lipid Z-Density Maps (left) and Contact Occupancy Structures (right) for all 12 lipid classes in the mammalian membrane.** Fig. S1 shows lipid species (LPC), glycosphingolipid (GM), ceramide (CER), phosphoinositide (PIP), phosphatidic acid (PA) and phosphotidylinositol (PI). Fig. S2 shows phosphatidylserine (PS), sphingomyelin (SM), phosphatidylethanolamine (PE), phosphatidylcholine (PC), cholesterol (CL) and diacylglycerol (DG). Density maps for each lipid type are show on the right. Contact occupancy structures are visualized from the intracellular view, the VSD-IV side view, the extracellular view and the VSD-II side view (left to right).

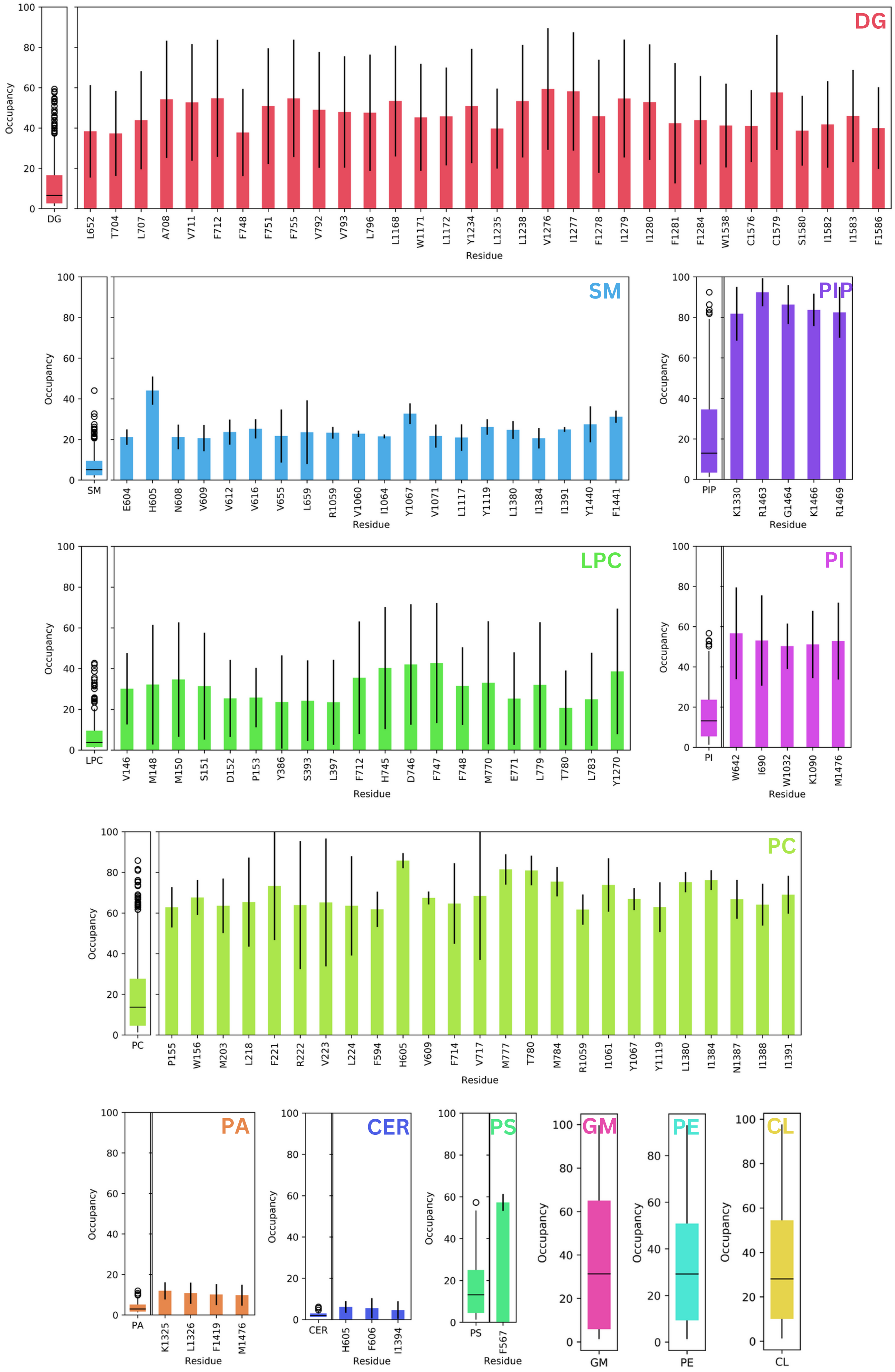


**Fig. S3 – Contact occupancy distributions (left) and outlying residues (right) for all 12 lipid classes in the mammalian membrane.**

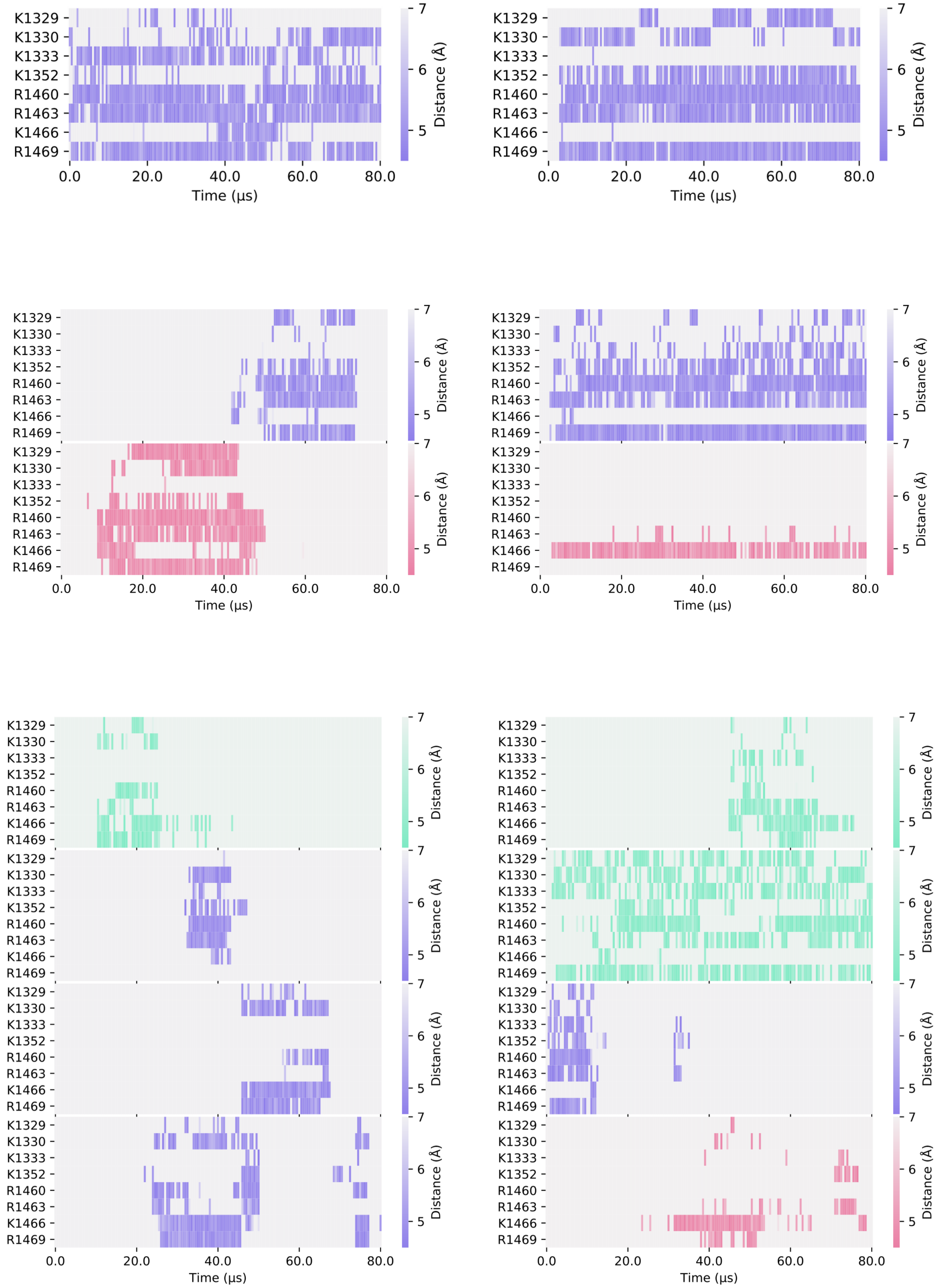
K1329
K1330
K1333
K1352
R1460
R1463
K1466
R1469
0.0
20.0
40.0
60.0
80.0
Time (μs)
Distance (Å)
5
6
7

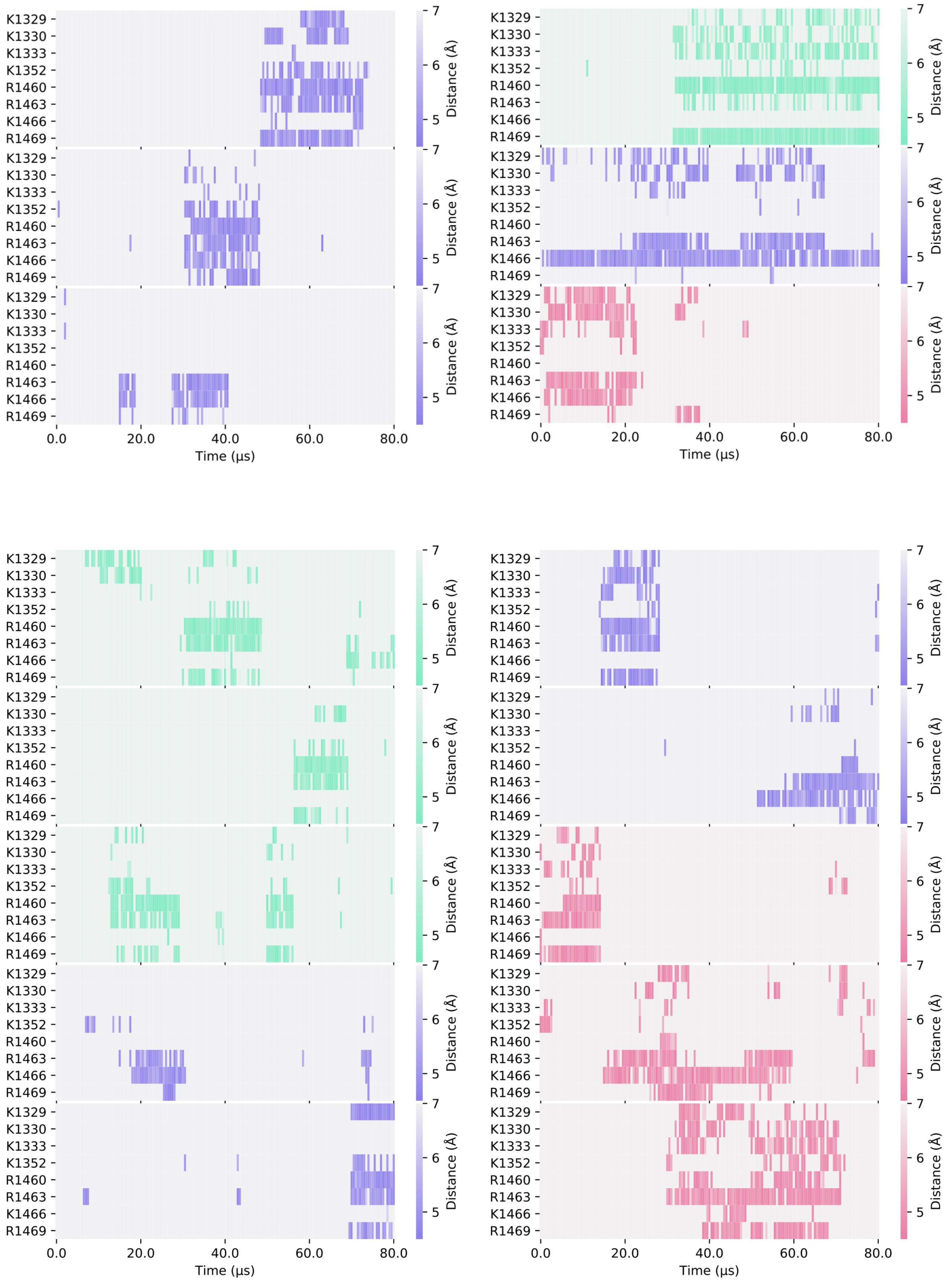


**Fig. S4-5 –Minimum distance between binding residues on $Na_v1.4$ and bound PIPs lipid across 31 long duration binding events (>10 μs), colored by distance and the type of PIP bound: PIP1 (blue-green), PIP2 (purple) and PIP3 (pink).**

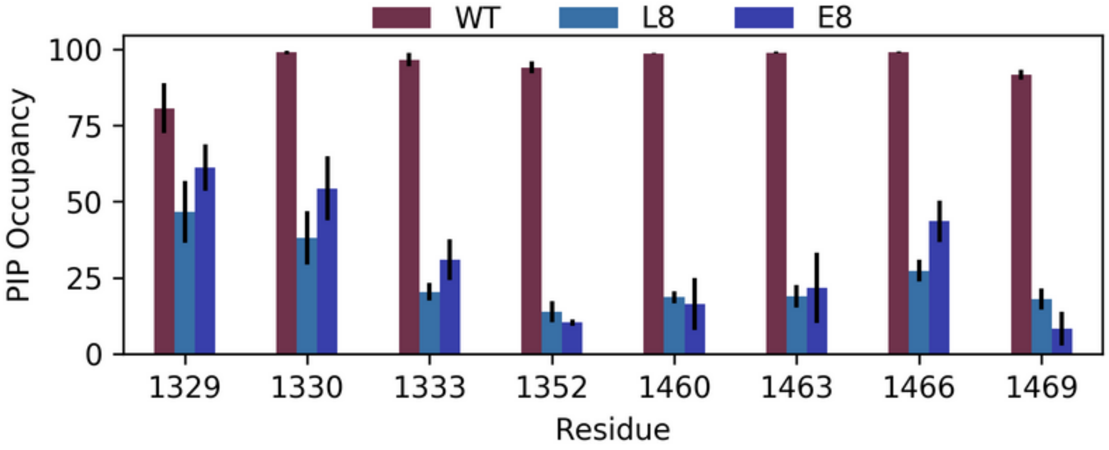


**Fig. S6 – PIP occupancy at putative binding site residues in WT (brown) and when all eight binding site resiudes are mutated to leucine (L8, light blue) or glutamate (E8, dark blue).**

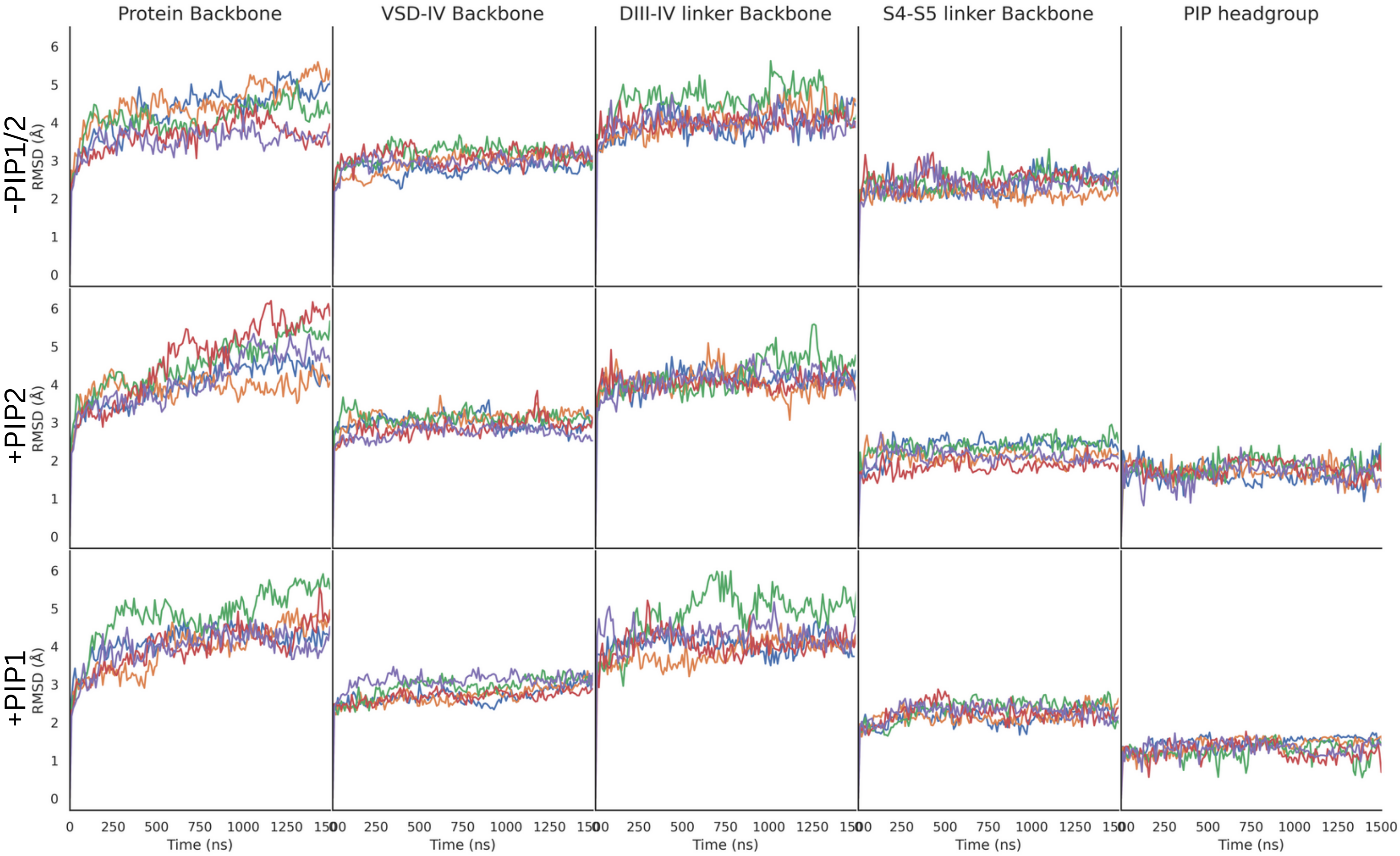


**Fig. S7 – Root Mean Square Deviation (RMSD) of the Na$_v$1.4 backbone, VSD-IV backbone, DIII-IV linker backbone and S4-S5 backbone (and PI(4,5)$P_2$/PI(4)$P_1$ headgroup) over 1.5 μs of atomistic simulations without any PIP (top row), with a single PI(4,5)$P_2$ lipid bound (middle row) and with PI(4)$P_1$ bound (bottom row) .** Five replicates were simulated – rep1 (blue), rep2 (red), rep3 (green), rep4 (orange), rep5 (purple).

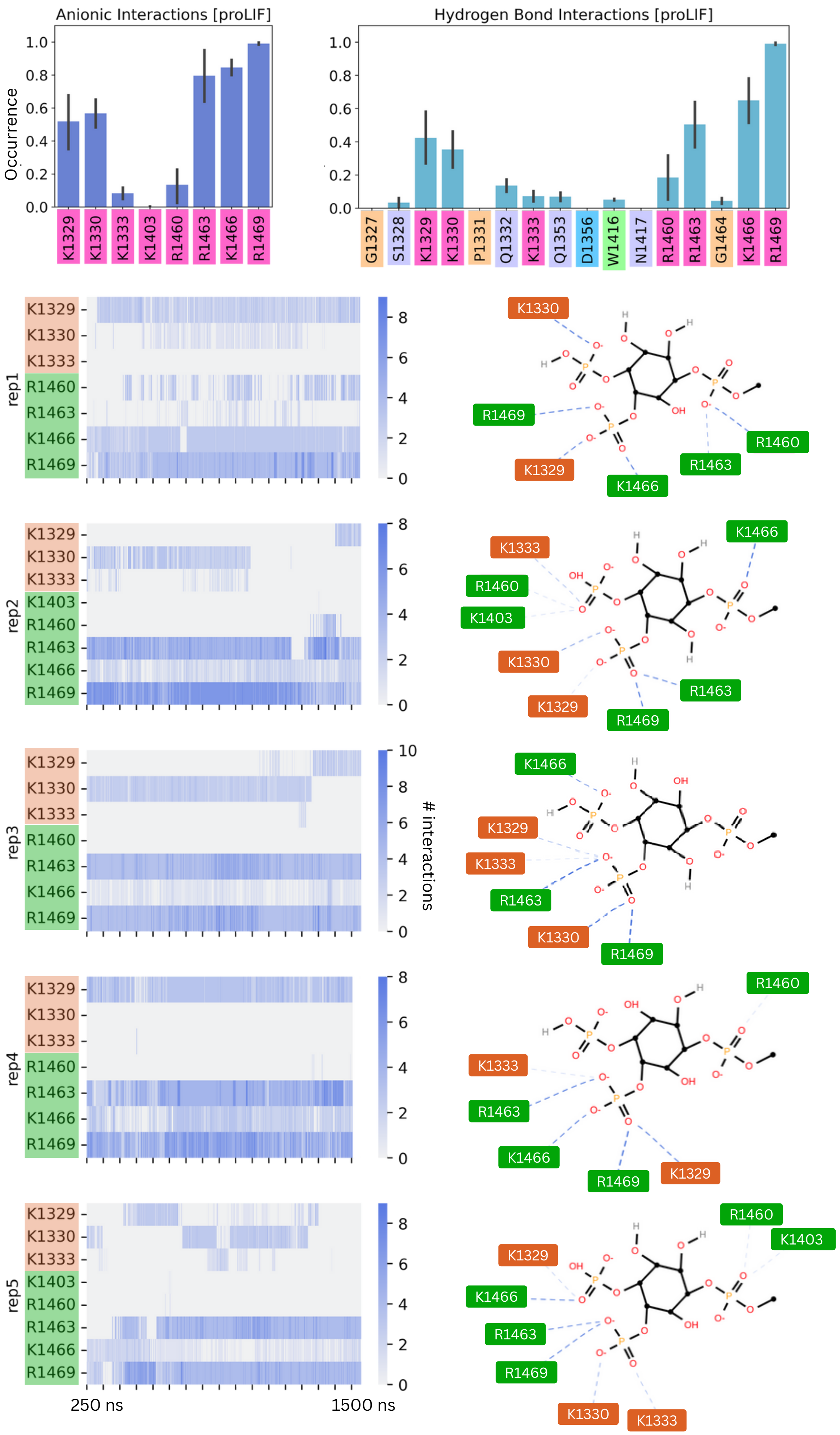


**Fig. S8 – Electrostatic interactions between the PI(4,5)$P_2$ headgroup and binding residues.** ProLIF bar charts describing the occurrence of anionic and hydrogen bond interactions between the PI(4,5)$P_2$ headgroup and binding residues (top). Per replicate data, showing the number of electrostatic interactions detected between the PI(4,5)$P_2$ headgroup and each of the binding residues (left); atomic interactions summarized in a ProLIF LigPlot (right).

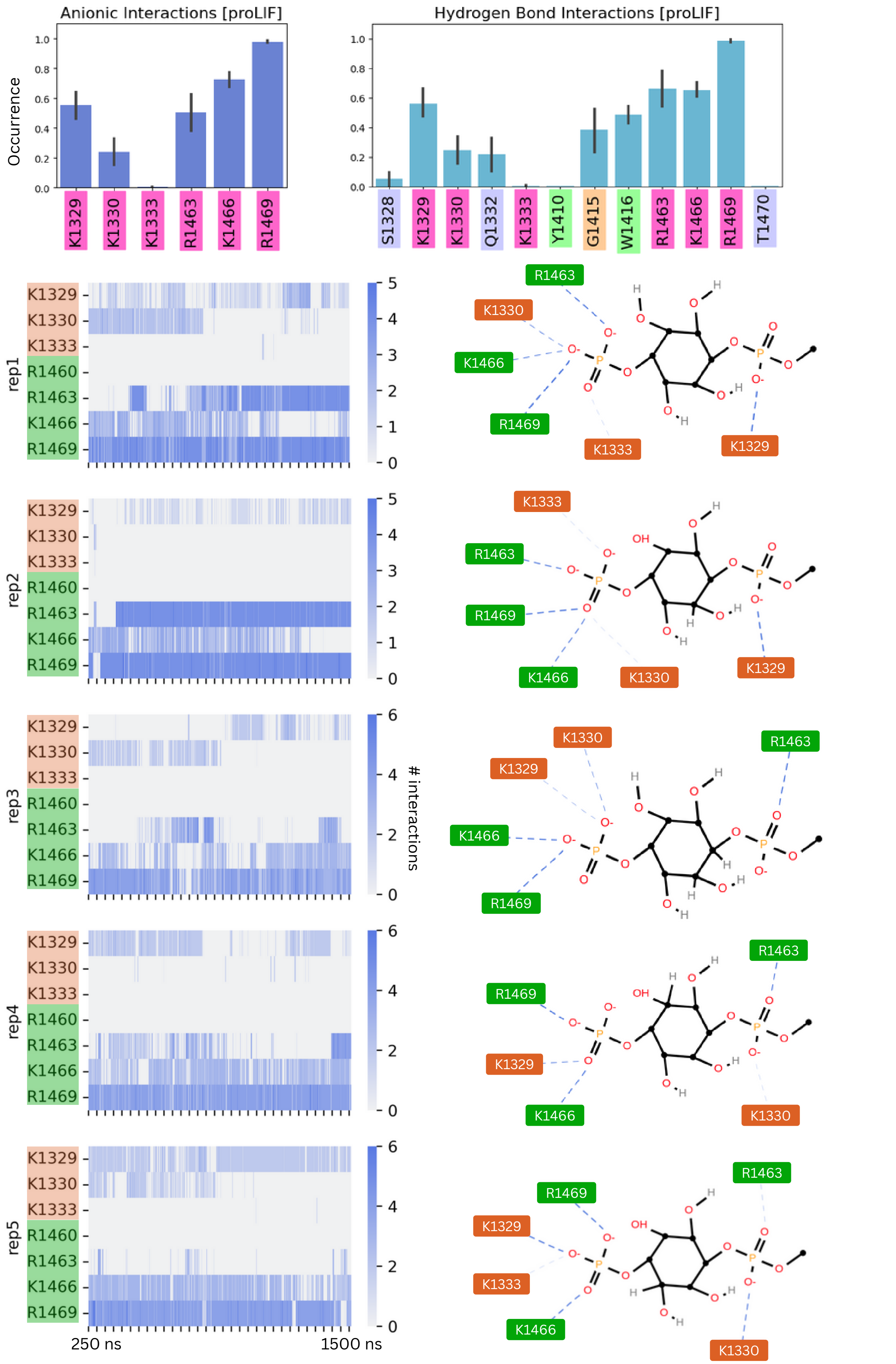


**Fig. S9 – Electrostatic interactions between the PI(4)$P_1$ headgroup and binding residues.** ProLIF bar charts describing the occurrence of anionic and hydrogen bond interactions between the PI(4)$P_1$ headgroup and binding residues (top). Per replicate data, showing the number of electrostatic interactions detected between the PI(4)$P_1$ headgroup and each of the binding residues (left); atomic interactions summarized in a ProLIF LigPlot (right).

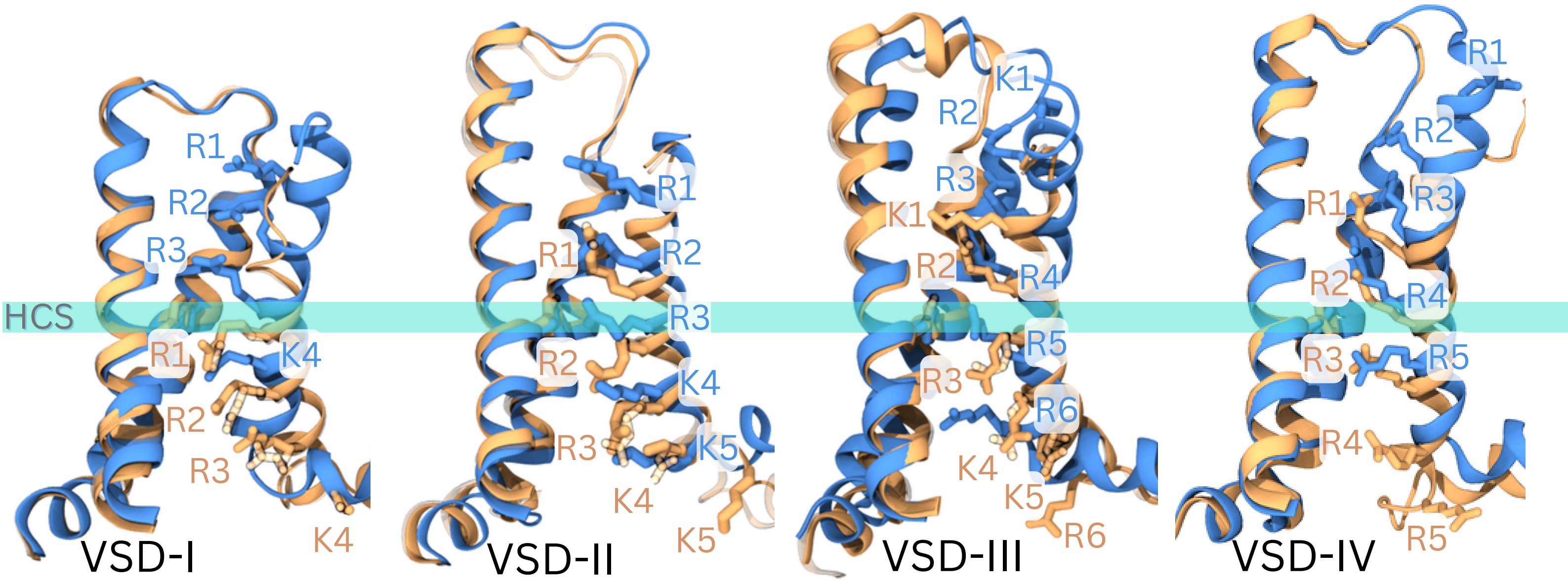


**Fig. S10 – Na$_v$1.7 resting state model validation.** Each of the four VSDs aligned separately, comparing the inactivated structure (6j8g, in blue) to the resting state model. All the S4 gating charges and the PHE/TYR residue on S2 to indicate the hydrophobic constriction site (HSC) for each VSD (in stick representation). The templates that each corresponding VSD was modelled from, is shown in translucent representation. S3 helix hidden for better visualisation of gating charge positions.

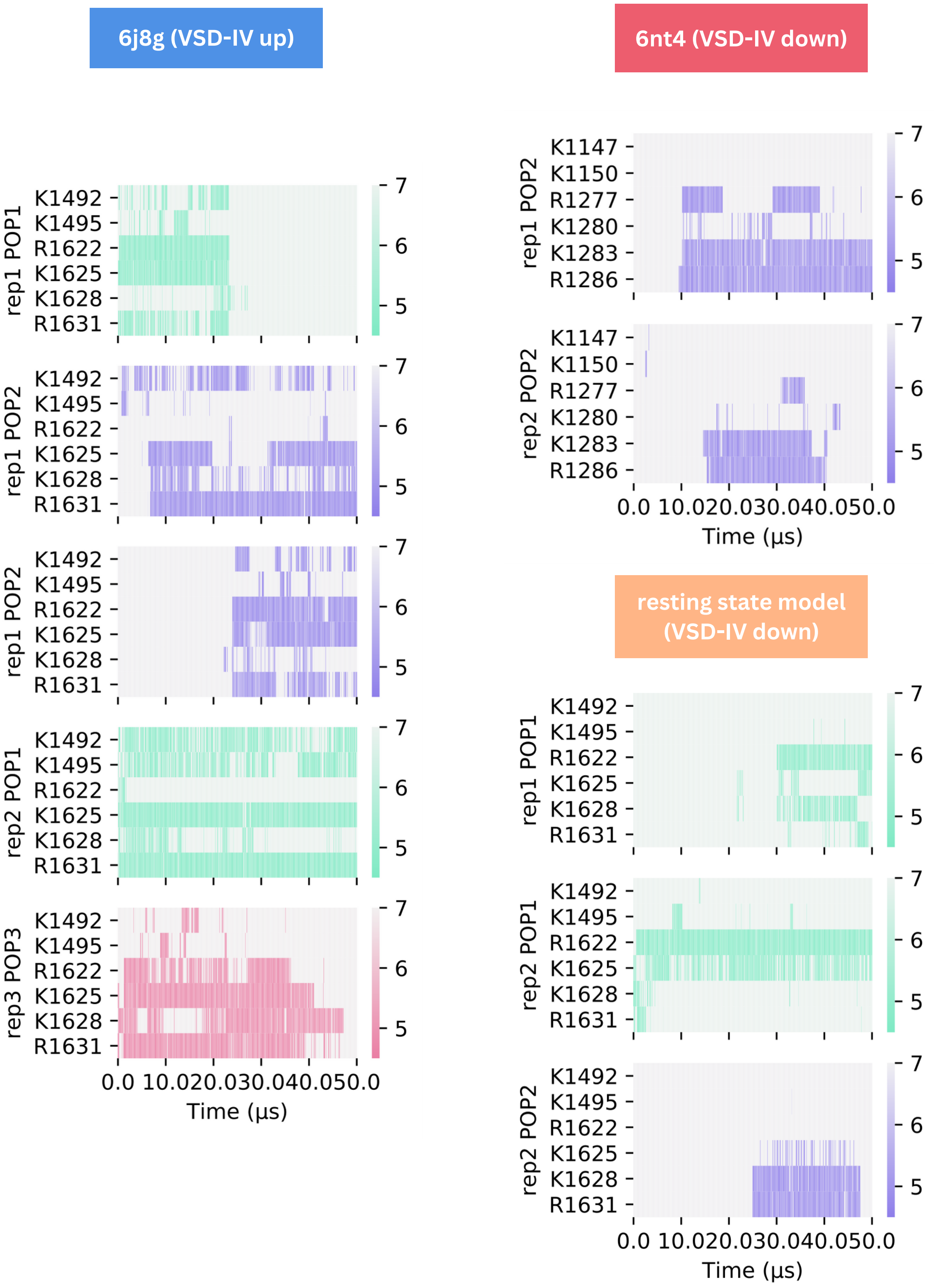


**Fig. S11 –Minimum distance between binding residues on $Na_v$1.7 and bound PIPs** shown for long duration binding events (>20 μs) in each system, colored by distance and the type of PIP bound: PIP1 (blue-green), PIP2 (purple) and PIP3 (pink). 5 long term binding events for the inactivated structure of $Na_v$1.7 (PDB ID: 6j8g) are shown on the left. $Na_v$1.7-$Na_v$Pas chimera (top right, PDB ID: 6nt4) and $Na_v$1.7 resting state model (bottom right) have fewer longer term interactions with PIP at this site. These interactions do not involve residues belonging to the DIII-IV linker (K1147/1492 and K1150/1495)

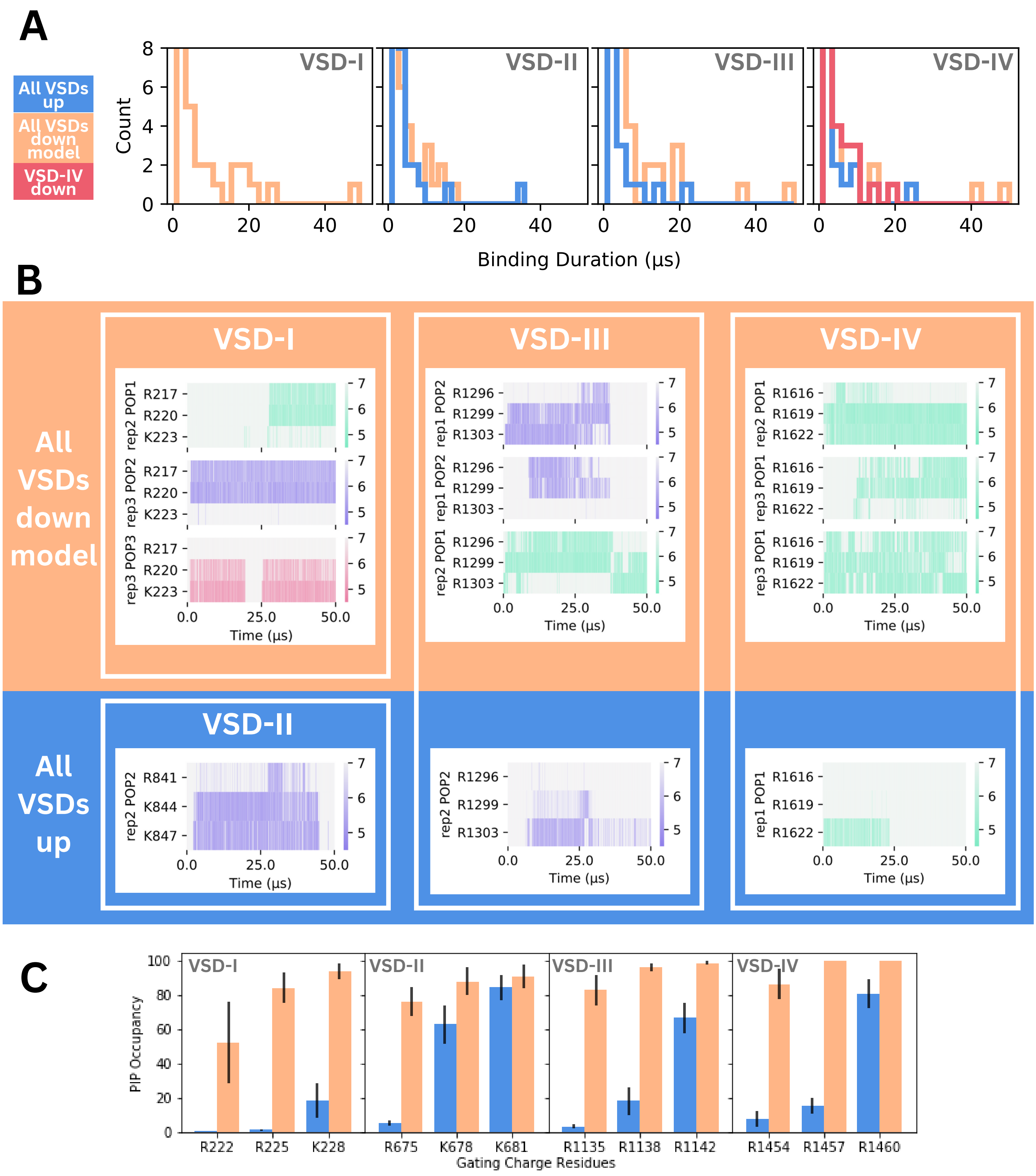


**Fig. S12 - PIP binding at S4 gating charges (A)** Binding duration distributions for PIP binding events at residues belonging to the bottom three gating charges in the S4 helix of **VSD I-III** are shown for the inactivated structure of Na$_v$1.7 (PDB ID: 6j8g, all VSDs up) and resting Na$_v$1.7 model (all VSDs down, CTD not modelled). For **VSD IV**, the distribution for the Na$_v$1.7-Na$_v$Pas chimera (VSD-IV down, CTD bound) is also shown. **(B)** Minimum distance between terminal three gating charges on each VSD and bound PIP for long duration (> 20 µs) binding events shown for the resting Na$_v$1.7 model (top) and inactivated Na$_v$1.7 structure (bottom). **(C)** Combined PIP occupancy at the bottom three gating charges on VSD I-IV in the inactivated (blue) and resting state model (orange) simulations of a Nav1.4 resting state model.
